# UHRF1 ubiquitin ligase activity supports the maintenance of low-density CpG methylation

**DOI:** 10.1101/2024.02.13.580169

**Authors:** Rochelle L. Tiedemann, Joel Hrit, Qian Du, Ashley K. Wiseman, Hope E. Eden, Bradley M. Dickson, Xiangqian Kong, Alison A. Chomiak, Robert M. Vaughan, Jakob M. Hebert, Yael David, Wanding Zhou, Stephen B. Baylin, Peter A. Jones, Susan J. Clark, Scott B. Rothbart

## Abstract

The RING E3 ubiquitin ligase UHRF1 is an established cofactor for DNA methylation inheritance. Nucleosomal engagement through histone and DNA interactions directs UHRF1 ubiquitin ligase activity toward lysines on histone H3 tails, creating binding sites for DNMT1 through ubiquitin interacting motifs (UIM1 and UIM2). Here, we profile contributions of UHRF1 and DNMT1 to genome-wide DNA methylation inheritance and dissect specific roles for ubiquitin signaling in this process. We reveal DNA methylation maintenance at low-density CpGs is vulnerable to disruption of UHRF1 ubiquitin ligase activity and DNMT1 ubiquitin reading activity through UIM1. Hypomethylation of low-density CpGs in this manner induces formation of partially methylated domains (PMD), a methylation signature observed across human cancers. Furthermore, disrupting DNMT1 UIM2 function abolishes DNA methylation maintenance. Collectively, we show DNMT1-dependent DNA methylation inheritance is a ubiquitin-regulated process and suggest a disrupted UHRF1-DNMT1 ubiquitin signaling axis contributes to the development of PMDs in human cancers.

## INTRODUCTION

Copying DNA methylation patterns onto newly synthesized DNA is a vital process in all dividing mammalian cells. Cytosines within CpG dinucleotides are the primary substrate for DNA methylation and are disproportionally distributed across the genome. Regions of high CpG density (i.e., CpG islands) account for ∼20% of all CpG sites and primarily occur in and around gene promoters and regulatory elements, while regions of low CpG density frequently occur in intergenic and intronic regions^1,2^. In normal cells, CpG islands are largely unmethylated, while the remaining ∼80% of CpGs throughout the genome are methylated^3,4^. An epigenetic hallmark of nearly all human cancers is a reversal of this normal DNA methylation pattern, where CpG island promoters acquire DNA hypermethylation associated with gene silencing of tumor suppressor genes (TSGs), while the remainder of the genome becomes hypomethylated^5–7^.

Despite a longstanding appreciation for intergenic DNA hypomethylation in cancer, the cause, function, and disease relevance of this epigenetic hallmark is not known. These DNA methylation losses induce formation of Partially Methylated Domains (PMDs) that occur almost exclusively in repressive and transcriptionally silent ‘B’ and ‘I’ compartments^8,9^. These higher-order chromatin compartments are characterized by repressive histone post-translational modifications (PTMs) like lysine 9 tri-methylation and lysine 27 tri-methylation on histone H3 (H3K9me3 and H3K27me3)^8–10^, nuclear lamina association^7^, late replication timing^11^, and low CpG density^12^. PMDs emerge in differentiated cells, are present across tissue types, and deepen in cancer cells and aging fibroblasts that undergo mitotic divisions^9–11,13–16^. Understanding how PMDs form is key to understanding the consequences of PMDs in cancer.

DNA methylation patterns are established early in development by the *de novo* methyltransferases DNMT3A and DNMT3B and are primarily maintained in pluripotent and somatic cells by DNMT1. DNMTs rely on histone and non-histone protein interactions to facilitate their epigenetic regulatory functions. This is exemplified by the role of UHRF1 (ubiquitin like with PHD and RING finger domains 1) in support of DNMT1-dependent maintenance methylation^17–19^. The RING E3 ubiquitin ligase UHRF1 is a multivalent epigenetic reader and writer that engages with chromatin via histone and DNA interactions^20^. The linked tandem Tudor (TTD) and plant homeodomain (PHD) finger of UHRF1 binds histone H3 through interactions with H3K9me2/me3 and the first 4 amino acids in the H3 N-terminal tail, respectively^21,23,25,27^. These histone binding domains also interact with DNA ligase 1 (LIG1) and SNF2 DNA helicase LSH to indirectly recruit UHRF1 to chromatin^22,24,26^. Engagement of H3K9me2/me3 enhances UHRF1 binding to hemi-methylated DNA (a DNA replication intermediate) through its SET- and RING-associated (SRA) domain^28,29^, and these interactions are necessary to direct its ubiquitin ligase activity towards H3K18, and to a lesser extent H3K14 and H3K23, on H3 tails^30,31^. These sites of mono-ubiquitination are binding sites for DNMT1 through tandem Ubiquitin Interacting Motifs (UIMs) embedded in its replication foci targeting domain (RFTS)^32–35^. Once recruited to chromatin, DNMT1 processively transfers methyl groups from S-adenosyl-methionine (SAM) to hemi-methylated CpG dinucleotide substrates, copying parent strand DNA methylation patterns to the newly replicated daughter strand^36–39^. While the link between UHRF1-mediated ubiquitination and DNMT1 chromatin targeting is established, the extent to which DNMT1 requires ubiquitin signaling for maintenance DNA methylation remains unclear.

In this study, we profiled the individual contributions of UHRF1 and DNMT1 to the maintenance of CpG methylation genome-wide using parallel doxycycline (dox)-inducible shRNA expression systems. We show that low-density CpGs are most prone to DNA hypomethylation in the absence of UHRF1. This loss of low-CpG density methylation contributes to the accelerated formation and deepening of PMDs and a reshaping of the DNA methylome that can be reversed with reintroduction of DNMT1, but not UHRF1. With genetic complementation studies of wild-type and domain loss-of-function UHRF1 and DNMT1 mutants, we show that DNMT1-dependent DNA methylation maintenance, independent of CpG density, requires ubiquitin reading, and that maintenance methylation at low-density CpG sites relies specifically on UHRF1 E3 ubiquitin ligase activity. Collectively, these studies demonstrate that DNA methylation maintenance is a ubiquitin-regulated process involving (but not exclusive to) UHRF1 enzymatic activity and suggest that a disrupted UHRF1-DNMT1 ubiquitin signaling axis contributes to the development of PMDs in human cancers.

## RESULTS

### Low-density CpGs are most prone to DNA hypomethylation when UHRF1 levels are reduced

To enable the comparative study of DNA methylation maintenance dynamics, we engineered HCT116 human colorectal carcinoma cells to express dox-inducible shRNAs targeting the *UHRF1* or *DNMT1* 3’UTRs (**Figure 1A, S1A**). High-resolution melt (HRM) analysis showed that UHRF1 depletion caused progressive loss of DNA methylation at promoters (*SFRP1)*, gene bodies (*STC2*), and pericentric heterochromatin (Chr6) over 2 weeks of Dox treatment (**Figure S1B**), while Dox treatment alone had no effect (**Figure S1C**). Using the Illumina EPIC array platform^40^, we profiled the individual contributions of UHRF1 and DNMT1 to DNA methylation maintenance at ∼800,000 individual CpGs from bulk and clonal cell populations at intervals up to two weeks following dox-inducible knockdown (**Figure 1A-B, S1A**). Clear differences in DNA hypomethylation dynamics were observed comparing UHRF1 and DNMT1 knockdown. Notably, DNMT1 knockdown caused hypomethylation of almost all highly methylated CpGs (β-value_Baseline_ ≥ 0.85), while UHRF1 knockdown revealed a sub-population of highly methylated CpGs that were resistant to DNA hypomethylation (**Figure 1B-C**).

**Figure 1.**
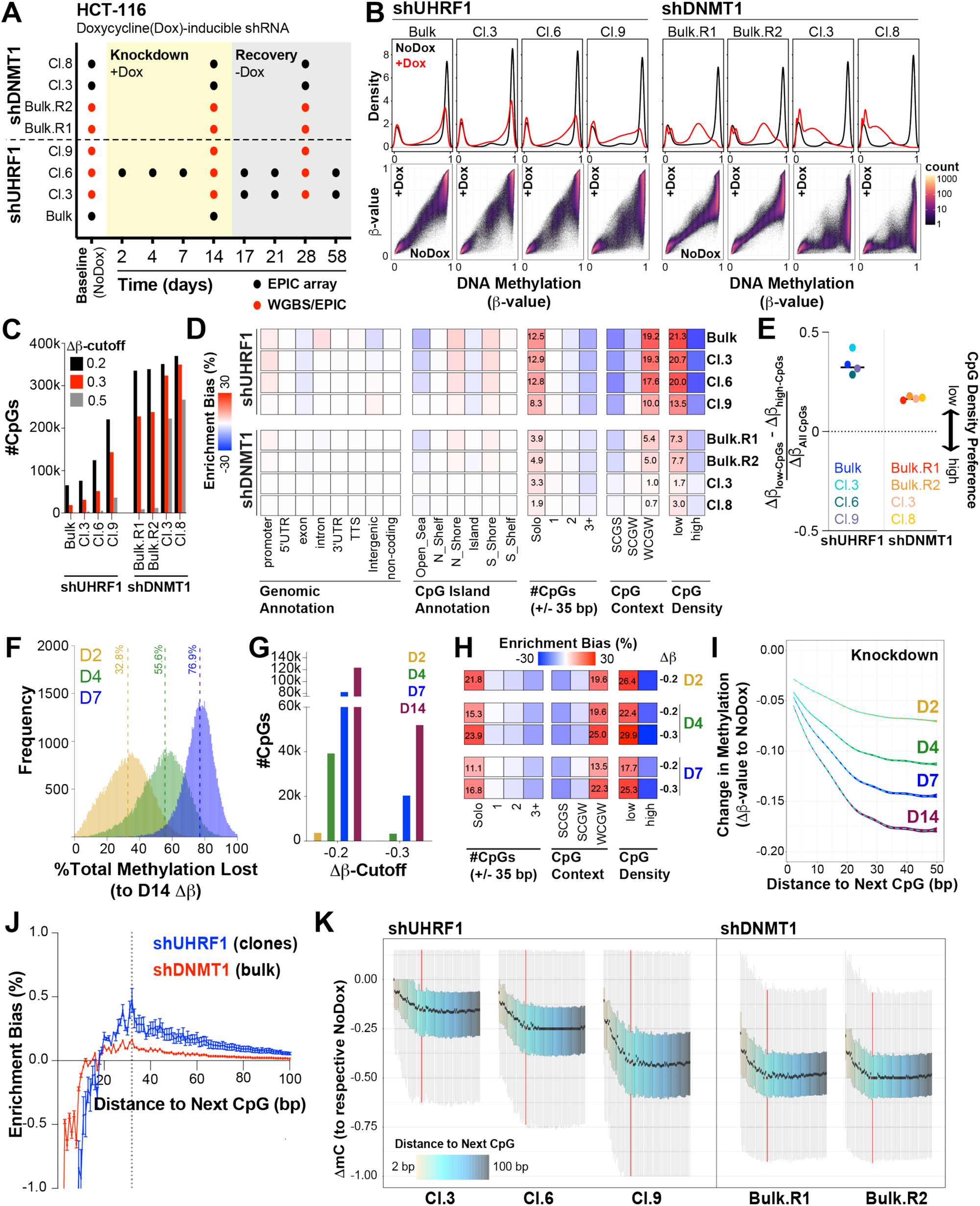
Low-density CpGs are most prone to DNA hypomethylation when UHRF1 levels are reduced. **(A)** Schematic of experimental design including samples, time-points, and types of DNA methylation data collected. **(B)** DNA methylation distributions of EPIC array probes from doxycycline (dox)-inducible shRNA HCT116 cell lines without (black) and with (red) dox treatment (10 ng/mL) for 14 days. **Top panel:** Density plots for CpG probe distribution across DNA methylation levels (β-value: 0 (unmethylated) to 1 (methylated)). **Bottom panel:** Density scatterplots demonstrating density of probes and DNA methylation level in baseline (x-axis) and knockdown (y-axis) methylomes. **(C)** Bar graph for number of differentially methylated CpGs (EPIC array) for each knockdown experiment (β-value_Baseline_ ≥ 0.85; varying Δβ-value_Knockdown_ cutoffs relative to each respective Baseline sample). **(D)** Hypergeometric analysis of significantly hypomethylated CpGs for each sample from **1C.** Red indicates significant overrepresentation for hypomethylation of the feature and blue indicates significant underrepresentation. Enrichment bias values are provided for the most significant positive enrichments. #CpGs indicates the number of CpGs ± 35 bp upstream and downstream of the hypomethylated CpG. CpG Context represents the −1/+1 position nucleotide (S = C or G; W = A or T) flanking the hypomethylated CpG. CpG density is determined by the number of bps to the next CpG (either upstream or downstream). Low density ≥ 20 bp, High Density < 20 bp. **(E)** Normalized preference for CpG density among the shUHRF1 and shDNMT1 samples. “CpG Density Preference” is calculated by subtracting the average Δβ-value of all the high-density CpGs (EPIC) from the average Δβ-value of the low-density CpGs, divided by the total Δβ-value of all CpGs. Positive values indicate preference for low-density CpGs and negative values preference for high-density CpGs. **(F)** Histogram of the calculated percent methylation lost ((β-value_Knockdown(X)_ – β-value_Baseline_)/Δβ-value_Knockdown(Day14)_)*100 across early time points (X = Day 2, 4, 7) for all significantly hypomethylated CpGs in shUHRF1 knockdown (Cl.6) from **1C**. Median % Methylation Lost is indicated by the dotted line for each time point. **(G)** Bar graph for number of differentially methylated CpGs (β-value_Baseline_≥ 0.85, varying Δβ-value_Knockdown_ cutoffs relative to Baseline) across the early time-points in shUHRF1 knockdown (Cl.6). **(H)** Hypergeometric analysis of significantly hypomethylated CpGs (from **1G**) across the early shUHRF1 (Cl.6) time-points. Legend from **1D** applies. **(I)** Average loss of DNA methylation for all highly methylated CpGs (β-value_Baseline_ ≥ 0.85) across ‘Distance to the Next CpG’ binning. Dotted line indicates the average Δβ-value as a function of distance, colored boundaries indicate 95% confidence intervals. **(J)** Hypergeometric analysis of significantly hypomethylated CpGs (WGBS) for each sample across CpG binning by ‘Distance to the Next CpG’ (bp). Positive enrichment bias indicates overrepresentation, negative enrichment bias indicates underrepresentation. Dotted line indicates peak positive enrichment bias for all shUHRF1 clones at 32 bp. **(K)** Boxplots for change in methylation (ΔmC_Knockdown_) of all highly methylated CpGs (mC_Baseline_ ≥ 0.85) from WGBS of each indicated sample across CpG binning by ‘Distance to the Next CpG’ (color bar [2-100 bp]). The 32 bp boxplot (peak enrichment bias from **1J**) is highlighted in red. **See also Figure S1.**

We next asked which CpGs were most prone to DNA hypomethylation (Δβ-value_Knockdown_ ≤ −0.3) in the absence of UHRF1 or DNMT1. To do this, we classified CpGs by locality (e.g., in genes or regulatory regions, CpG island status) and by CpG density. Two relative CpG density measurements were used. The first CpG density measure is where the number of adjacent CpGs (+/− 35 bp) and flanking nucleotide sequences (S = C/G, W = A/T) are jointly considered (i.e. ‘Solo-WCGW’)^11^. The second defines “high-density” CpGs as the distance between two adjacent CpGs being < 20 bp apart (upstream or downstream), and “low-density” CpGs are those where CpGs are ≥ 20 bp apart. We then performed enrichment bias analysis (normalized to the distribution of CpGs with β-value_Baseline_ ≥ 0.85), and we found that CpG density (by both density measurements) was the primary attribute that defined DNA hypomethylation in the absence of UHRF1 (**Figure 1D**). While DNMT1 knockdowns demonstrated a slight enrichment bias towards low-density CpGs (**Figure 1D**), normalization of low-to high-density CpG hypomethylation showed that UHRF1 knockdowns consistently had a higher preference for low-density CpG hypomethylation (**Figure 1E**). These findings were verified by whole genome bisulfite sequencing (WGBS)^41^ (**Figure S1D,E**). These results are consistent with a prior study linking ‘Solo-WCGW’ hypomethylation to UHRF1 depletion^42^.

To further investigate the relationship between UHRF1 and CpG density, we next asked if UHRF1 preferentially bound CpGs in the ‘Solo-WCGW’ context. Fluorescence polarization binding assays showed that UHRF1 did not have a ‘WCGW’ sequence bias for binding to hemi-methylated CpGs (**Figure S1F**). We then considered if the enrichment bias for ‘Solo-WCGW’ could be attributed to an inherent bias present in the human genome for low-density CpGs and A/T flanking nucleotide sequences. Indeed, enrichment bias analysis of all CpGs in the genome showed that ‘WCGW’ CpGs are inherently low-density CpGs, as the likelihood of being a ‘WCGW’ CpG increases with decreasing CpG density (**Figure S1G,H**). Finally, we asked if the average methylation level of low-density CpGs was equivalent to ‘Solo-WCGW’ CpGs across 291 cell lines previously profiled on the Illumina 450K array^43^. These data showed that the average β-value of low-density CpGs is highly correlated with the ‘Solo-WCGW’ CpG average DNA methylation level but incorporates over 100,000 additional probes into the analysis (**Figure S1K**). Collectively, these data show that maintenance DNA methylation at low-density CpGs depends on UHRF1, independent of sequence context.

To extend this analysis, we next profiled DNA methylation in a clonally expanded, Dox-inducible, shUHRF1 cell population (Cl.6) at different time-points following UHRF1 knockdown (**Figure 1A, F-I**). Consistent with HRM analysis of candidate loci (**Figure S1B**), deepening hypomethylation of CpGs occurred over two weeks of UHRF1 knockdown (**Figure 1F**), especially at low-density CpGs (**Figure 1G,H**). Importantly, the degree of DNA hypomethylation (Δβ-value_Knockdown_) again was dependent on CpG density (**Figure 1I**). Given this result, we next asked whether there was an enriched peak distance to the next CpG among the hypomethylated CpGs (from **Figure S1D**). We found an inflection of peak distance enrichment for CpG hypomethylation at 32 base pairs for the shUHRF1 knockdown clones, an enrichment that was attenuated by DNMT1 knockdown (**Figure 1J**). Profiling of all highly methylated CpGs (mC_Baseline_ ≥ 0.85) by WGBS likewise showed the greatest DNA hypomethylation at low-density CpGs, with a plateau of loss at ∼32 base pairs (highlighted in red; **Figure 1K**). Notably, mathematical modeling of DNA methylation maintenance kinetics (derived from Repli-BS-seq^44^) predicted that DNMT1 acts processively on CpGs that are up to 36 base pairs apart^45^. Collectively, these data suggest that maintenance DNA methylation mechanisms are not conserved throughout the genome and that perhaps CpG density influences how DNA methylation is maintained.

### Hypomethylation of low-density CpGs reshapes the DNA methylome in the absence of UHRF1 and DNMT1

Hypomethylation at low-density CpG sites is linked to the cellular mitotic index and the formation of PMDs, especially in late replicating chromatin^11,15^. Indeed, it has been hypothesized that the increased mitotic index in cancer shortens the available time for cells to complete methylation maintenance before the next cell division, resulting in passive hypomethylation in late replicating genomic regions. Our data are consistent with this hypothesis, in that HCT116 colorectal carcinoma cells are hypomethylated at low-density CpGs in late replicating genomic regions at baseline, while high- and low-density CpGs in early replicating chromatin are hypermethylated (**Figure 2Ai, Figure S2A,Bi**).

**Figure 2.**
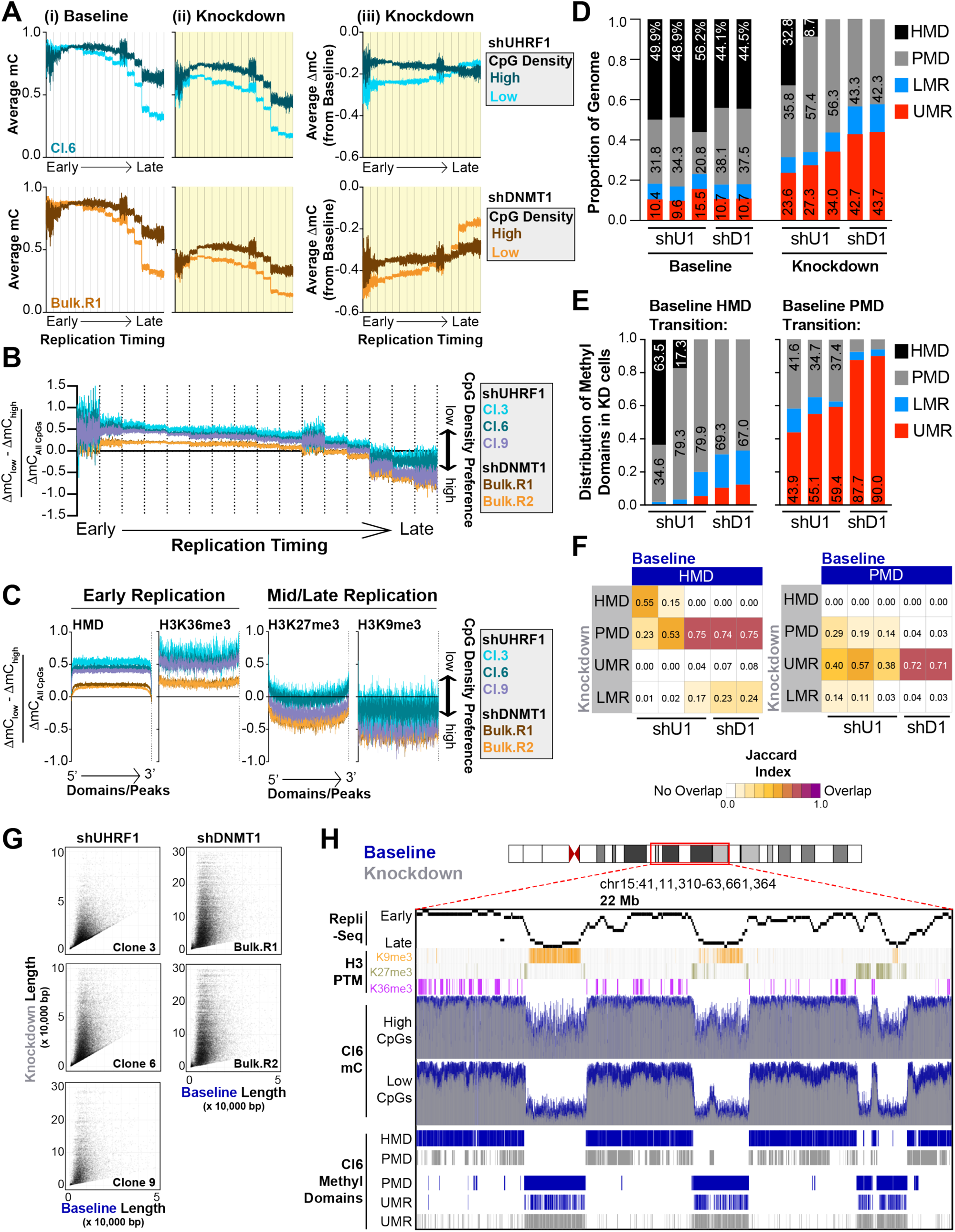
Hypomethylation of low-density CpGs in the absence of UHRF1 and DNMT1 reshapes the DNA methylome. **(A)** Average DNA methylation (mC) of high (dark color) and low (light color) density CpGs in (i) Baseline and (ii) Knockdown phases and the (iii) average change in methylation (ΔmC_Knockdown_) across each replication timing phase (16-phase) in HCT116 cells. Replication timing phases are assigned in 50 kb windows genome-wide, and the average DNA methylation (from WGBS) is calculated in 100 bp bins from the start of the 50 kb window to the end for each timing phase. Vertical lines indicate the separation of the different replication timing phases from early to late. shUHRF1 Cl.6 and shDNMT1 Bulk (replicate 1) samples are provided in the main figure. Remaining WGBS samples are presented in **Figure S2A. (B)** Normalized preference for CpG density in shUHRF1 and shDNMT1 samples across replicating timing phases. ‘CpG Density Preference’ is calculated by subtracting the average ΔmC of all the high-density CpGs (WGBS) from the average ΔmC of the low-density CpGs, divided by the total ΔmC of all CpGs in 100 bp bins as described in **2A**. Positive values indicate preference for low-density CpGs and negative values for high-density CpGs. **(C)** Normalized preference for CpG density in shUHRF1 and shDNMT1 samples across genomic features known to be localized in early replicating chromatin (Highly Methylated Domains (HMDs), H3K36me3) and mid/late replicating domains (H3K27me3/H3K9me3). DNA methylation is averaged from the 5’ end of the Domain/Peak to the 3’end in size normalized windows. **(D)** Proportional coverage of the genome for called Methylation Domains in the Baseline and Knockdown stage for each shUHRF1 (order: Cl.3,6,9) and shDNMT1 (order: Bulk R1, R2) WGBS sample. HMD = Highly Methylated Domain, PMD = Partially Methylated Domain, LMR = Lowly Methylated Region, UMR = Unmethylated Region. Baseline mC distributions for Methylation Domains are provided in **Figure S2C**. **(E)** Distribution shift of called Methylation Domains in the Knockdown methylome from Baseline HMDs (left) and PMDs (right). Order of samples is same as Figure 1D. **(F)** Overlap analysis of called Methylation Domains from Baseline and Knockdown methylomes across shUHRF1 and shDNMT1 WGBS samples. The Jaccard Index measures the extent of the overlap with 0 being no overlap to 1 being complete overlap. **(G)** Scatterplot of UMR length in Baseline methylomes versus Knockdown methylomes. Over 90% of the UMRs located within Baseline PMDs expand with loss of UHRF1 and DNMT1. **(H)** Browser shot of 22 Mb region on Chr15 demonstrating the observations made from WGBS sequencing analysis of Baseline (blue) and Knockdown (gray) methylomes in the shUHRF1 Cl.6 sample integrated with Repli-Seq, histone PTMs, and locality of called Methylation Domains. **See also Figure S2.**

Given the preferential hypomethylation of low-density CpGs when UHRF1 is absent, we next asked how depleting UHRF1 or DNMT1 affected DNA methylation at high- and low-density CpGs in early- and late-replicating chromatin. In early replicating chromatin, depleting UHRF1 or DNMT1 induced hypomethylation at high- and low-density CpGs, but the extent of hypomethylation was deeper for low-density CpGs (**Figure 2Aiii, Figure S2Biii**). UHRF1 knockdowns demonstrated a stronger preference than DNMT1 knockdowns for hypomethylation at low-density CpGs (**Figure 2B**), similar to what we observed by EPIC array analysis (**Figure 1E**). In late-replicating chromatin, low-density CpGs had a lower DNA methylation level at baseline than high-density CpGs (**Figure 2Ai**), and both UHRF1 and DNMT1 knockdowns drove low-density CpGs to become almost completely demethylated (**Figure 2Aii**). As low-density CpGs became deeply hypomethylated (**Figure 2Aii**), high-density CpGs in late-replicating chromatin became the preferential target for hypomethylation in the absence of UHRF1 or DNMT1 (**Figure 2Aiii,B**). As an orthogonal analysis, we profiled the CpG density preference for hypomethylation in the absence of UHRF1 or DNMT1 using annotations of histone PTMs and methylation domains known to localize with early (highly methylated domains (HMDs), H3K36me3) and mid/late replicating chromatin (H3K27me3/H3K9me3) **(Figure 2C**)^8,9^. Indeed, HMDs and H3K36me3-marked chromatin demonstrated increased preference for hypomethylation of low-density CpGs in UHRF1 knockdowns over DNMT1 knockdowns, while H3K27me3/H3K9me3-marked chromatin did not (**Figure 2C**).

As hypomethylation at low-density CpG sites generates PMDs^11,12,15^, we next asked how DNA methylation domains throughout the genome were altered with UHRF1 or DNMT1 knockdown (**Figure S2C**). For both UHRF1 and DNMT1 knockdown, we found substantial restructuring of the DNA methylome where HMDs were lost, and expanded coverage of the genome by PMDs and unmethylated regions (UMRs) emerged (**Figure 2D**). HMDs predominately transitioned into PMDs (**Figure 2E,F**) through preferential hypomethylation at low-density CpGs (**Figure 2C**), while pre-existing UMRs (present within PMDs) expanded in size (**Figure 2E-G**). Deep hypomethylation adjacent to the original UMR (**Figure S2D**) initiated UMR expansion, with loss of DNA methylation at both high- and low-density CpGs (**Figure S2E**). Collectively, these data show that depleting UHRF1 or DNMT1 substantially reshapes the DNA methylome (**Figure 2H**), forming PMDs in early-replicating chromatin and expanding UMRs in late-replicating chromatin.

### Recovery of low-density CpG methylation requires UHRF1

Above, we used the dox-inducible knockdown system to understand how DNA methylation patterns change when UHRF1 or DNMT1 are knocked down. Here, we used the same system to understand how the DNA methylome recovers from a hypomethylated state when UHRF1 or DNMT1 are re-expressed (**Figure 1A**). In this case, we treated the engineered HCT116 human colorectal carcinoma cells with dox for two weeks (to generate hypomethylated genomes), terminated the dox treatment, and then monitored DNA methylation recovery for two weeks as UHRF1 or DNMT1 expression was restored (**Figure S3A**).

To profile DNA methylation recovery of the most significantly hypomethylated CpGs (from analysis in **Figure S1D**), we compared changes in DNA methylation from the end of the recovery phase (28 days) to the original baseline DNA methylation measurements for each individual CpG (ΔmC_Recovery_). The DNA methylation recovery measurements were then binned into three groups, where CpGs with recovery values (|ΔmC_Recovery_|*100) < 10% are considered not recovered, > 90% are considered complete recovery, and between 10 and 90% are considered incomplete or partial recovery (**Figure S3B**). Following re-expression of DNMT1, > 85% of the hypomethylated CpGs were re-methylated to within 90% of their baseline value (**Figure 3A, B**). Recovery was almost universal across CpGs of varying densities, although the minority of CpGs that did not recover their baseline methylome were enriched for low-density CpGs (**Figure S3C**). We obtained similar results using EPIC arrays and clonally expanded, dox-inducible DNMT1 knockdown lines (**Figure S3D-F**). After UHRF1 re-expression, however, CpG re-methylation was incomplete (**Figure 3A,B**). Enrichment bias analysis showed that high-density CpGs had near complete recovery, but that low-density CpGs had partial to incomplete recovery (**Figure 3C**). Consistent with these data, profiling of all baseline highly methylated CpGs (β-value_Baseline_ ≥ 0.85) demonstrated that DNA methylation recovery was more complete for high-density CpGs while low-density CpGs remained the most demethylated with re-introduction of UHRF1 (**Figure 3D**). Indeed, low-density CpG methylation never returned to the baseline state, even after 58 days of restored UHRF1 expression (**Figure S3G-H**).

**Figure 3.**
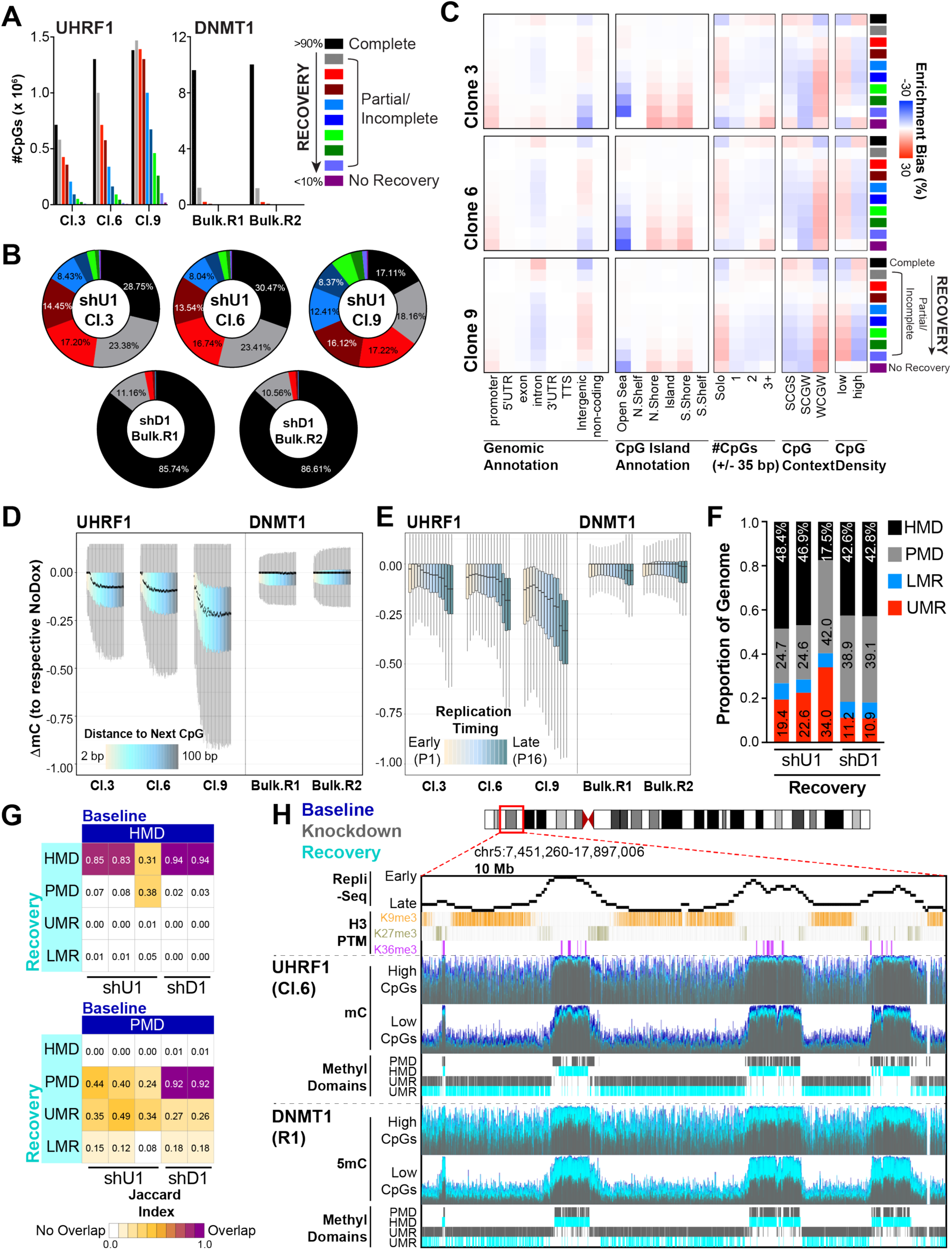
Recovery of low-density CpG methylation requires UHRF1. **(A)** Recovery analysis for significantly hypomethylated CpGs (from **Figure S1D**) across shUHRF1 and shDNMT1 (WGBS) samples. DNA methylation recovery is determined by calculating the ΔmC of Day 28 samples (Recovery) to the respective Baseline (NoDox) samples. DNA methylation recovery measurements were then binned into three groups, where CpGs with recovery values (|ΔmC_Recovery_|*100) < 10% are considered not recovered, > 90% are considered complete recovery, and between 10 and 90% are considered incomplete or partial recovery. **(B)** Distribution of recovery categories among shUHRF1 and shDNMT1 samples (WGBS). Scale from **3A** applies. **(C)** Hypergeometric analysis of DNA methylation recovery for each shUHRF1 sample (WGBS) from **3A.** Red indicates significant overrepresentation for recovery of the feature and blue indicates significant underrepresentation. **(D)** Boxplots for change in methylation recovery (ΔmC_Recovery_) of all highly methylated CpGs (mC_Baseline_ ≥ 0.85) from WGBS of each indicated sample across CpG binning by ‘Distance to the Next CpG’ (color bar [2-100 bp]). **(E)** Boxplots for change in methylation recovery (ΔmC_Recovery_) of all highly methylated CpGs (mC_Baseline_ ≥ 0.85) from WGBS of each indicated sample across replication timing phases. **(F)** Proportional coverage of the genome of called Methylation Domains in the Recovery stage for each shUHRF1 and shDNMT1 sample (WGBS). **(G)** Overlap analysis of called Methylation Domains from Baseline and Recovery methylomes across shUHRF1 and shDNMT1 WGBS samples. The Jaccard Index measures the extent of the overlap with 0 being no overlap and 1 being complete overlap. **(H)** Browser shot of 10 Mb region on Chr5 demonstrating the observations made from WGBS analysis of the Recovery (light blue) methylome in shUHRF1 Cl.6 and shDNMT1 replicate 1 samples integrated with Repli-Seq, histone PTMs, and locality of called Methylation Domains. **See also Figure S3.**

We next considered how replication timing integrated with DNA methylation recovery dynamics. While DNMT1 re-introduction universally recovered DNA methylation across replication timing phases, UHRF1 re-introduction primarily restored DNA methylation levels in early replicating chromatin but not late replicating chromatin (**Figure 3E**). UHRF1 re-expression also restored HMDs in early replicating regions of the genome (particularly in Cl.3 and Cl.6), but the expanded UMRs never reverted to their original state (**Figure 3F-H**). Collectively, these data show that HCT116 cells can fully restore their DNA methylation patterns after DNMT1 loss and re-expression, but that late replicating low-density CpG methylation patterns are not restored after UHRF1 loss and re-expression. These data suggest that UHRF1 creates an epigenetic memory or ‘bookmark’ that DNMT1 uses to re-establish DNA methylation patterns at low-density CpGs.

### UHRF1 ubiquitin ligase activity is required for maintenance of low-density CpG methylation

The appreciated substrates for UHRF1 enzymatic activity are mono-ubiquitination sites on histone H3 tails and PAF15^31,32,46–48^. We therefore hypothesized that the ‘bookmark’ required for DNMT1 to mediate recovery of low-density CpG methylation was UHRF1-dependent ubiquitination. To test this hypothesis, wild-type (WT) and various domain loss-of-function mutant UHRF1 transgenes were stably introduced into the dox-inducible UHRF1 shRNA line that showed the deepest loss in UHRF1 expression and DNA demethylation in the knockdown state (Cl.9; **Figure 4A, S4A**). UHRF1 transgene expression in the absence of dox treatment had no effect on DNA methylation (**Fig S4B**). Consistent with prior reports^49–51^, a mutation in the aromatic cage of the UHRF1 TTD that disrupts binding to H3K9me2/me3 (Y188A) had no effect on maintenance DNA methylation, while a SRA mutation that abolishes DNA binding (G448D) had the same effect as dox-inducible UHRF1 knockdown cells complemented with an empty vector control (EV) (**Figure 4A-B**). In contrast, UBL (F46V and F59V)^52,53^ or RING (H741A)^54^ mutations that perturb ubiquitin transfer had partial effects on DNA methylation maintenance (**Figure 4A-B, S4C**). We then used enrichment bias analysis to identify the subset of CpGs dependent on UHRF1 ubiquitin ligase activity for DNA methylation maintenance. UHRF1 ubiquitin ligase activity was needed to support DNA methylation maintenance in low CpG density regions of the genome in both HCT116 and RKO colon cancer cell lines (**Figure 4C-E, S4D-F**), consistent with histone ubiquitination functioning as a bookmark for low-density CpG methylation. Collectively, these data support the hypothesis that UHRF1 provides a ubiquitin ‘bookmark’ that is essential for the maintenance of low-density CpG methylation.

**Figure 4.**
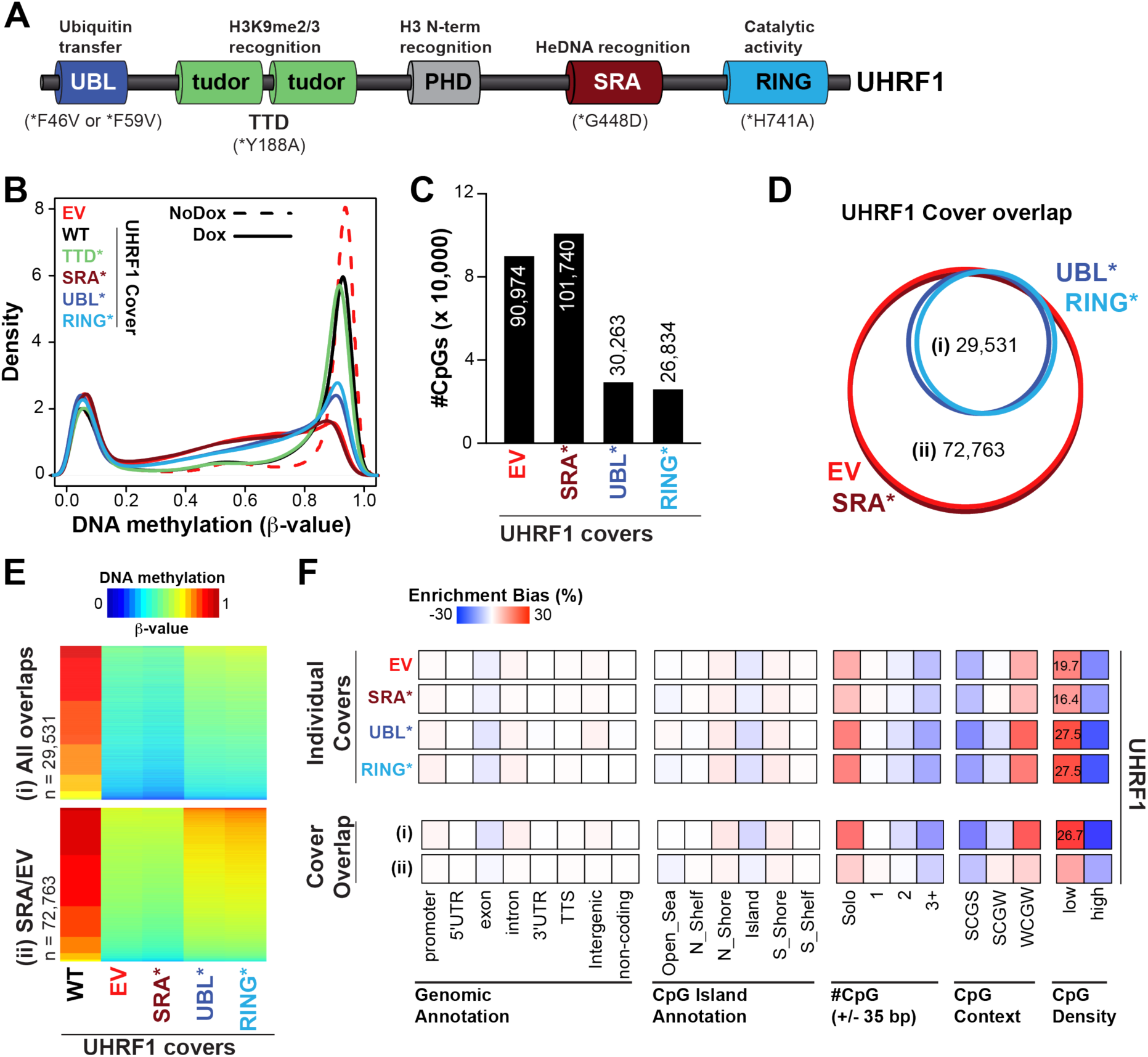
UHRF1 ubiquitin ligase activity is required for the maintenance of low-density CpG methylation. **(A)** Schematic of UHRF1 protein domains and mutations made to the UHRF1 transgene covers. **(B)** Density plot for CpG probe distribution across DNA methylation levels (β-value: 0 (unmethylated) to 1 (methylated)) for WT and mutant(*) UHRF1 transgene covers. **(C)** Bar graph of number of hypomethylated CpGs (EPIC array) for each UHRF1 cover (β-value_WT_cover_ ≥ 0.85; Δβ-value_mutant_covers_ ≤ −0.3). **(D)** Venn diagram demonstrating overlap of hypomethylated CpGs from UHRF1 cover experiments. **(E)** Heatmap of DNA methylation (β-value) values for overlapping hypomethylated CpGs from the (i) Ubiquitin mutant covers and (ii) EV and SRA mutant covers. **(F)** Hypergeometric analysis of hypomethylated CpGs from UHRF1 cover experiments. Red indicates significant overrepresentation for hypomethylation of the feature (Genomic annotation, CpG island annotation, etc.) and blue indicates significant underrepresentation. Enrichment bias values are provided for the most significant positive enrichments. **See also Figure S4.**

### DNMT1 ubiquitin recognition is essential for DNA methylation maintenance

To further test the hypothesis that ubiquitin serves as a ‘bookmark’ for DNMT1-mediated DNA methylation maintenance, we next considered the contributions of the DNMT1 UIMs to its DNA methylation maintenance function (**Figure 5A**). *In vitro* methyltransferase activity assays with recombinant full-length, WT and domain loss-of-function DNMT1 mutants (**Figure 5A**) showed that UIM mutations had no effect on the catalytic activity of DNMT1 toward free DNA (**Figure 5B**). However, *in* vitro binding assays with recombinant DNMT1 RFTS domain showed that RFTS binding to H3K14/18/23ub nucleosomes required UIM2 and partially depended on UIM1 (**Figure 5C, S5A**), suggesting that the two UIMs make separate and unequal contributions to DNMT1 recruitment to ubiquitinated nucleosomes. Mutations to both UIMs (dblUIM) completely abolished DNMT1 RFTS binding to ubiquitinated nucleosomes.

**Figure 5.**
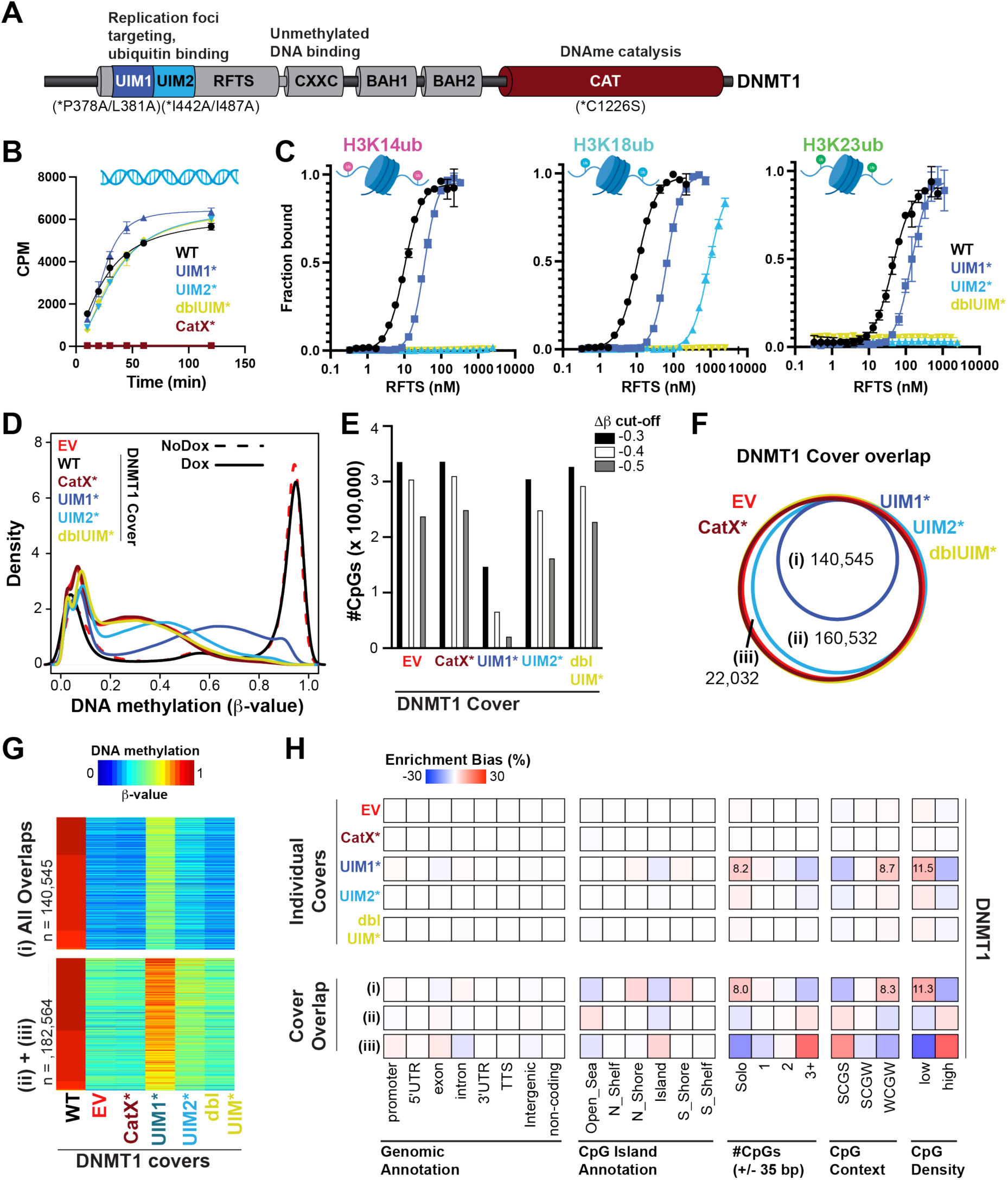
DNMT1 ubiquitin recognition is essential for DNA methylation maintenance. **(A)** Schematic of DNMT1 protein domains and mutations made to the DNMT1 transgene covers. **(B)** *In vitro* methyltransferase assays measuring catalytic activity of recombinant full length DNMT1 wild type (WT), Catalytic Dead (CatX), and UIM mutants toward free hemi-methylated DNA. **(C)** *In vitro* Alpha-screen proximity assays measuring interactions between H3 ubiquitinated nucleosomes (K14ub, K18ub, K23ub) and recombinant DNMT1 RFTS domain wild type (WT) and UIM mutants. **(D)** Density plot for CpG probe distribution across DNA methylation levels (β-value: 0 (unmethylated) to 1 (methylated)) for WT and mutant(*) DNMT1 transgene covers. **(E)** Bar graph of number of hypomethylated CpGs (EPIC array) for each DNMT1 cover (β-value_WT_cover_ ≥ 0.85; Δβ-value_mutant_covers_ ≤ −0.3, −0.4, −0.5). **(F)** Venn diagram demonstrating overlap of hypomethylated CpGs from DNMT1 cover experiments. **(G)** Heatmap of DNA methylation (β-value) values for overlapping hypomethylated CpGs from (i) All overlaps and (ii+iii) UIM2, dblUIM, CatX, and EV overlaps. **(H)** Hypergeometric analysis of hypomethylated CpGs from DNMT1 cover experiments. Red indicates significant overrepresentation for hypomethylation of the feature and blue indicates significant underrepresentation. Enrichment bias values are provided for the most significant positive enrichments. **See also Figure S5.**

Next, we used a genetic complementation strategy (as in **Figure 4**) to determine how UIM mutants contribute to DNA methylation maintenance (**Figure 5A, S5B)**. DNMT1 transgene expression in the absence of dox treatment had no effect on DNA methylation (**Figure S5B-C**), and a WT DNMT1 transgene cover maintained nearly all DNA methylation in the absence of endogenous protein (**Figure 5D**). Consistent with perturbed binding to ubiquitinated nucleosomes (**Figure 5C, S5A**), mutations to UIM2 (both individual and in dblUIM) abolished the ability of DNMT1 to maintain DNA methylation much like a catalytically dead enzyme (CatX; **Figure 5D,E**). As the DNMT1 UIM mutants maintain catalytic activity *in vitro* (**Figure 5B**), these results collectively indicate that DNMT1 requires recruitment to chromatin through ubiquitin interaction to perform DNA methylation maintenance. UIM1 mutation, however, did not completely abolish maintenance methylation (**Figure 5D,E**). Rather, DNA methylation maintenance was attenuated (**Figure 5D,E**) much like the ability of DNMT1 RFTS with UIM mutation to bind ubiquitinated nucleosomes (**Figure 5C).** Indeed, the DNMT1 UIM1 mutant was still able to maintain DNA methylation at > 50% of all highly methylated CpGs (β-value_WT_ ≥ 0.85) while mutants in UIM2 and the catalytic domain could not (**Figure 5F,G**). Maintenance DNA methylation in the absence of UIM1 was most notably affected in regions of low CpG density (**Figure 5H**), consistent with a role for UHRF1 ubiquitin ligase activity promoting DNMT1 function specifically through UIM1. Taken together, these data demonstrate that ubiquitin reading activity is essential for DNMT1-mediated DNA maintenance methylation and that UHRF1 ubiquitin ligase-dependent recruitment of DNMT1 through UIM1 is necessary to support DNA methylation in regions of the genome with low-density CpGs. These data also suggest that additional E3 ligases function redundantly with or independent of UHRF1 to support DNA methylation maintenance elsewhere in the genome.

## DISCUSSION

In this study, we showed that UHRF1 ubiquitin ligase activity and DNMT1 ubiquitin reader function are essential for the maintenance of low-density CpG methylation. While mutations to the RING and UBL domains of UHRF1 perturbed maintenance methylation of low-density CpGs, these mutants could maintain methylation patterns at higher-density CpGs. Conversely, mutation of the SRA domain (which abolishes the interaction of UHRF1 with DNA) disrupted maintenance DNA methylation of additional CpGs, suggesting that UHRF1 aids DNMT1 in the maintenance of DNA methylation through both ubiquitin-dependent and independent mechanisms. Furthermore, it is clear from the growing body of co-factors (LIG1^22^, LSH^24,26^) and additional substrates for UHRF1-directed ubiquitination (PAF15^47,48^, H3^31,32^), that maintenance DNA methylation is not a conserved process genome-wide but rather a coordinated effort among different mechanisms to ensure the faithful inheritance of DNA methylation prior to cell division.

In support of this notion, profiling of DNA methylation maintenance kinetics (via coupling BrdU/EdU labelling of newly replicated DNA with next-generation sequencing^42,44,55^ and mass spectrometry^56^) has made clear that maintenance methylation does not solely occur in the immediate wake of the replication fork, but rather can be divided into replication-fork coupled (rapid) and replication-fork uncoupled (delayed) phases. Additionally, the kinetics of DNA methylation maintenance are not conserved across CpG dinucleotides, as high-density and low-density CpGs demonstrate rapid and delayed methylation rates, respectively^42,44,45,57^. Indeed, mathematical modeling of maintenance DNA methylation kinetics (from Repli-BS data^44^) emphasized the importance of CpG density with regard to maintenance methylation rates^45^. Remarkably, this modeling predicted that DNMT1 acts rapidly and processively on CpGs that are, on average, within 36 base pairs of a neighboring CpG^45^.

The field has converged on the importance of CpG density for maintenance DNA methylation, particularly within a window of 32-36 bps to the next CpG, in several additional ways. First, the ‘solo-WCGW’ hypomethylation signature observed in PMDs defines ‘solo’ CpGs as those lacking additional CpGs within 35 bp upstream or downstream of its location^11^. Second, depletion of UHRF1 enriches for hypomethylated CpGs that are ∼32 bp from neighboring CpGs (**Figure 1J**). Third, the degree by which CpGs are hypomethylated in response to loss of both UHRF1 and DNMT1 deepens with increasing distance to neighboring CpGs but plateaus at ∼35 bps (**Figure 1K**). Collectively, these data suggest that DNMT1 acts processively if it encounters the next CpG within ∼35 bp. After ∼35 bp, DNMT1 requires additional recruitment mechanisms to ensure fidelity of low-density CpG maintenance methylation. We hypothesize that UHRF1 ubiquitin ligase activity functions as a compensatory mechanism for the lack of DNMT1 processivity in regions with low CpG density.

### Connecting PMD formation to the maintenance DNA methylation machinery

The implications of low-density CpG hypomethylation can be observed from methylomes of aged and cancerous tissues where PMDs deepen due to loss of DNA methylation maintenance at these CpGs^9,11,58–60^. The ‘solo-WCGW’ signature was first described within PMDs where CpGs in this context are most prone to DNA methylation loss^11^, and we and others have linked UHRF1 depletion to hypomethylation of these CpGs^42^. Our WGBS analysis of the methylome in the absence of UHRF1 revealed that hypomethylation of low-density CpGs that reside in baseline HMDs results in the formation of PMDs. To our knowledge, this is the first study to report a direct perturbation to a DNA methylation writer or co-factor that induces formation of PMDs. To date, PMD formation through loss of low-density CpG methylation has largely been associated with the mitotic index of the cells as PMDs deepen through progressive cell divisions^15^. However, the exact mechanism that contributes to PMD formation remains unknown. We hypothesize that UHRF1 dysfunction (specifically to the ubiquitination axis) contributes to the formation of PMDs in ageing and cancer. Importantly, we observe a divergence in the ability of the methylome to recover with re-introduction of UHRF1 as early replicating regions regain their HMD status while late replicating regions with deepened PMDs do not recover. This observation supports the paradigm that the fidelity of DNA methylation maintenance is dependent on replication timing where early replicating chromatin (with both high- and low-density CpGs) is faithfully copied while late replicating chromatin (enriched in low-density CpGs) has limited time to ensure this process is complete before mitosis.

### Disruption of UHRF1 ubiquitin ligase activity is compatible with tumorigenesis

We previously showed that UHRF1 supports the growth and metastasis of human colorectal cancer (CRC) through repression of TSG expression^49^. Notably, UHRF1 functional domains do not contribute equally to CRC maintenance. Histone and DNA interactions mediated by the PHD and SRA domains, respectively, are essential for repression of TSGs and oncogenic growth. However, perturbations to the RING domain that induce hypomethylation of low-density CpGs (**Figure S4F**) are dispensable for oncogenic growth and proliferation^49^, suggesting that loss of low-density CpG methylation is compatible with tumorigenesis. Indeed, PMDs (through hypomethylation of low-density CpGs) are universally observed across cancer methylomes^11^. Our results suggest that the erosion of DNA methylation during oncogenesis (at low-density CpGs) is connected to dysregulated UHRF1 ubiquitin ligase signaling.

### CpG density is a potential substrate specificity determinant for UHRF1 enzymatic activity

In addition to H3 tails, UHRF1 also mono-ubiquitinates the PCNA associated factor PAF15 to facilitate an interaction with DNMT1^31,32,47,48^. It has been proposed that PAF15 ubiquitination serves as a DNMT1 recruitment mechanism in early replicating chromatin, whereas H3 tail ubiquitin recruits DNMT1 in late replication^48^. So how is the ubiquitin ligase activity of UHRF1 directed towards certain substrates for maintenance of low-density CpG methylation? *In vitro* ubiquitination assays measuring UHRF1 activity towards semi-synthetic nucleosome substrates provides a potential biochemical explanation for this association between CpG methylation density and UHRF1-dependent H3 ubiquitination^46,53^.

Specifically, increasing hemi-methylation density in linker DNA directs UHRF1 enzymatic activity away from histone H3 – presumably due to a geometric constraint associated with DNA binding through the SRA domain^46,53^. Considering these data with genome-level analyses presented in this paper, we hypothesize that CpG density controls the substrate specificity of UHRF1, and that H3 in genomic regions of high CpG density are poor substrates for UHRF1 ubiquitin ligase activity.

### DNMT1 and the role of ubiquitin signaling for DNA methylation maintenance

Differences in recovery dynamics following UHRF1 and DNMT1 knockdown were surprising to us. While DNMT1 re-introduction essentially restored DNA methylation patterns to the pre-knockdown state, UHRF1 re-introduction showed inefficient recovery of low-density CpG methylation in late replicating regions of the genome. This imbalance in low-density CpG methylation mirrors patterns of DNA hypomethylation reported in primary human cancers. We thus speculate that UHRF1-dependent H3 ubiquitination ‘bookmarks’ genomic regions of low CpG density in late replicating chromatin for re-methylation by DNMT1, and that a disrupted UHRF1-DNMT1 ubiquitin signaling axis contributes to DNA hypomethylation in cancer. This hypothesis is consistent with the model that links PMD formation in cancer to mitotic divisions^11,15^, where competition between the pace of DNA replication and cell division leads to incomplete maintenance DNA methylation. A requirement for signaling through a multi-step write-read-write mechanism of ubiquitin-dependent DNA methylation maintenance may be error prone and unresolvable late in the replication cycle.

Also surprising to us was the observation that ubiquitin reading activity, while dispensable for DNMT1 activity *in vitro*, is required to support the full extent of DNMT1-dependent DNA methylation maintenance in cells. DNMT1 adopts an autoinhibitory conformation when the RFTS (containing its UIMs) sits within the catalytic core^33,34,61–64^. Interaction with multi-monoubiquitinated H3 partially releases the RFTS (N-lobe, UIM1) from the catalytic pocket and enhances the methyltransferase activity of DNMT1^34,39,64^. Integrating this observation with our results, we suggest that ubiquitin signaling provides two primary functions for DNMT1-mediated maintenance methylation – partial relief from an autoinhibited state, and recruitment of DNMT1 to replicating chromatin. As disruptions to UHRF1 and its ubiquitin ligase activity only partially affect DNMT1-dependent DNA methylation, our data also suggests that other ubiquitin ligases, functioning in a compensatory or independent manner with UHRF1, contribute to DNMT1 maintenance methylation activity. Indeed, NEDD4, CUL4A, and PHF7 all have reported overlapping H3 ubiquitin substrate specificity with UHRF1^65–69^.

Finally, we report for the first time distinct functions of the two DNMT1 UIMs, both in binding ubiquitinated nucleosomes and in DNMT1 maintenance methylation activity. The convergent phenotypes of disrupting UHRF1 ubiquitin ligase activity and DNMT1 UIM1 ubiquitin reading activity suggest that DNMT1 UIM1 reads the ubiquitin ‘bookmark’ laid down by UHRF1 to maintain DNA methylation in regions of low CpG density. Loss of DNMT1 UIM2 reading activity, on the other hand, disrupted DNMT1 activity almost to the same extent as catalytic inactivation of the enzyme, suggesting that DNMT1 UIM2 supports DNA methylation maintenance genome-wide. DNMT1 UIM2 may function in concert with other ubiquitin ligases that target the H3 tail, and it may also have critical roles in maintenance methylation beyond the recognition of ubiquitin on the H3 tail.

### Concluding remarks

In summary, we have demonstrated that the ubiquitin ligase activity of UHRF1 is essential for supporting DNMT1-mediated maintenance methylation of low-density CpGs. Additionally, we have provided evidence that disruption to the UHRF1-DNMT1 ubiquitin signaling axis promotes low-density CpG hypomethylation patterning reminiscent of PMD formation observed in ageing and cancer. Finally, we have shown that DNMT1 requires interaction with ubiquitin to mediate DNA methylation maintenance. Collectively, these results demonstrate the complexity of maintaining DNA methylation patterns in dividing cells and provide crucial insight for future work aimed at understanding these mechanisms.

## ACKNOWLEDGEMENTS

We thank members of the Rothbart laboratory for helpful discussions. We also thank Benjamin Johnson for discussions on bioinformatic analysis and Darrell Chandler for critical reading of the manuscript.

Computation for the work described in this paper was supported by the High-Performance Cluster and Cloud Computing (HPC3) Resource at the Van Andel Institute. We thank the Van Andel Institute Genomics Core (RRID:SCR_022913), especially Marie Adams, for assistance with processing EPIC arrays and next-generation sequencing.

## Funding

The American Cancer Society Michigan Cancer Research Fund PF-16-245-01-DMC (R.L.T.)

National Institutes of Health R35GM124736 (S.B.R.)

National Institutes of Health P50CA254897 (S.B.R., S.B.B., P.A.J.)

The American Cancer Society RSG-21-031-01-DMC (S.B.R.)

National Health and Medical Research Council (NHMRC) Investigator Grant #1156408 (S.J.C)

NHMRC Ideas Grant 2012072 (S.J.C., Q.D)

NHMRC Investigator Grant #1177792 (Q.D.)

National Institutes of Health F32CA260116 (J.H.)

## AUTHOR CONTRIBUTIONS

Conceptualization: R.L.T., S.B.R.

Methodology: R.L.T., J.H., Q.D., S.J.C., S.B.R.

Software: R.L.T., Q.D.

Validation: R.L.T., J.H., Q.D., A.K.W., H.E.E., A.A.C.

Formal Analysis: R.L.T., J.H., Q.D.

Investigation: R.L.T., J.H., Q.D., A.K.W, H.E.E., B.M.D., X.K., A.A.C., R.M.V., W.Z.

Resources: R.L.T., S.B.R., J.M.H., Y.D.

Data Curation: R.L.T., Q.D.

Writing – Original Draft: R.L.T., S.B.R.

Writing – Review & Editing: All authors

Visualization: R.L.T, J.H., Q.D.

Supervision: S.B.R., S.J.C., P.A.J.

Project Administration: S.B.R.

Funding Acquisition: R.LT., S.B.R., J.H., P.A.J., S.B.B., S.J.C., Q.D.

## METHODS

### Generation of HCT116 cells with doxycycline-inducible shRNAs

#### Tet-pLKO-puro vector cloning

Empty Tet-pLKO-puro plasmid^70^ was purchased from Addgene (#21915), and shRNA oligos for insertion (**Table 1**) were synthesized by GenScript. Oligos were reconstituted in water at a concentration of 100 μM, diluted in 10X annealing buffer (1 M NaCl, 100 mM Tris, pH 7.4), brought to 100°C for 10 mins, and naturally cooled to 30°C to anneal the oligo pair. Annealed oligos were diluted 1:400 in 0.5X annealing buffer. Tet-puro-pLKO plasmid was digested with AgeI and EcoRI (NEB) restriction enzymes, gel-purified (NEB), and ligated to the annealed shRNA oligos. Ligated plasmids were transformed into stable competent *E. Coli* (High Efficiency) cells (NEB) and incubated overnight at 37°C on ampicillin LB agar plates. Clones were selected and grown in liquid LB culture with ampicillin, and the plasmid was purified by miniprep (Qiagen). Purified plasmids were screened for the shRNA oligo insert using XhoI digestion and sequenced for validation.

**Table 1.**
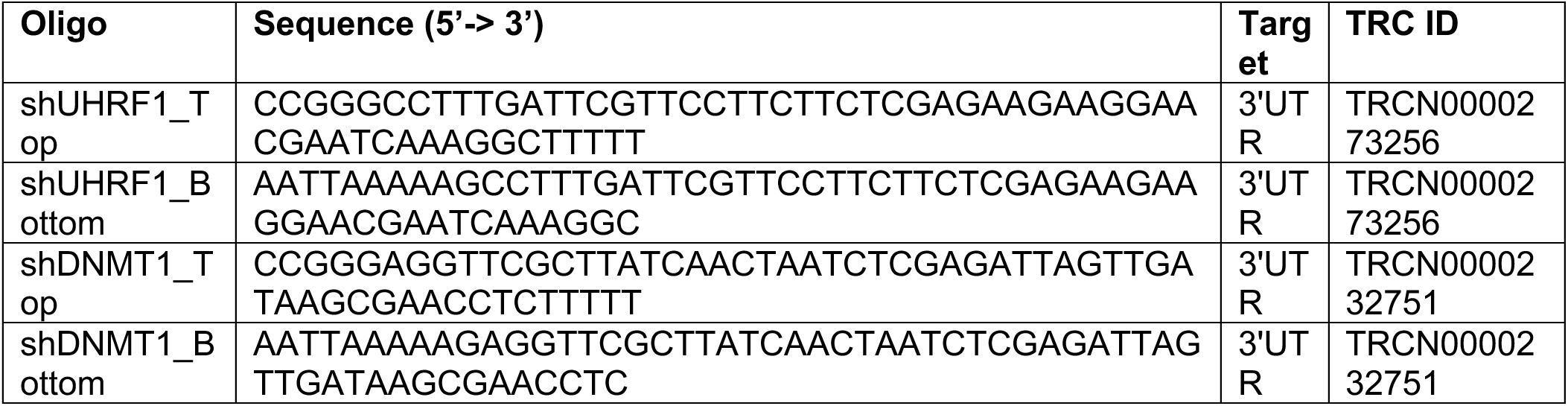
Oligo sequences for shRNA cloning.

#### Lentiviral production for dox-inducible shRNA integration

Lentivirus production for the stable integration of dox-inducible shRNAs was conducted in HEK 293T cells purchased from ATCC. HEK 293T cells were grown in DMEM supplemented with 10% FBS to ∼70% confluency on 60 mm cell culture plates prior to transfection at 37°C with 5% CO_2_. Cells were then transfected with the Tet-pLKO-puro plasmids targeting either *UHRF1* or *DNMT1* transcripts and the accompanying lentiviral packaging (psPAX2) and envelope (pMD.2G) plasmids with Opti-MEM and Xtreme HP Gene transfection reagent (Roche) per the manufacturer’s protocol. After 15 hours of incubation, media was refreshed for an additional 24 hours of incubation. Media containing lentiviral particles was then removed, collected, and stored at 4°C. Media was refreshed for a second 24-hour incubation, and then collected and pooled with the first viral media collection. Viral media was cleared of cell debris by centrifugation at 700 x g for 5 minutes followed by passage through a 0.45 micron filter (Avantor PES 25 mm 0.45 μm). Virus was aliquoted and stored at −80°C prior to transduction.

#### Transduction and clonal selection of HCT116 cells

HCT116 cells were purchased from ATCC and maintained in McCoy’s 5A Media with 10% FBS and 1% Penicillin/Streptomycin at 37°C with 5% CO_2_. HCT116 cells were plated in 6-well plates and infected with media containing 8 μg/mL polybrene (with no antibiotics) and 500 μl of shRNA lentivirus. Media was refreshed 24 hours post-infection, and the cells were allowed to grow for an additional 24 hours prior to puromycin selection (2 μg/mL) for 2 days. An uninfected plate of HCT116 cells was grown in parallel to test puromycin resistance in the infected cells. Clonal populations for both shUHRF1 and shDNMT1 HCT116 cell lines were isolated using serial dilution of 5000 cells in 96-well plates. Wells containing single cells were noted and allowed to propagate until a sufficient number of cells for freezing and maintenance were attained.

### Generation of HCT116 dox-inducible shRNA cell line with UHRF1/DNMT1 transgene covers

We selected HCT116 dox-inducible shUHRF1 and shDNMT1 clonal cell populations (Clone 9 and Clone 3, respectively) that demonstrated the deepest loss in DNA methylation with dox-inducible knockdown of the endogenous target protein. Selection of these clones allowed us to deplete endogenous protein with dox and query DNA methylation maintained in the presence of wild-type (WT) or mutant transgene protein covers.

#### Retrovirus production for UHRF1/DNMT1 wild-type and mutant transgene covers

UHRF1, DNMT1 and respective mutants (**Table 2**) were cloned into the pMXs-IRES-blasticidin retroviral vector (a gift from David Sabatini, Addgene #72876) by EcoRI and XhoI restriction sites without an affinity tag. Phoenix AMPHO cells (purchased from ATCC) were transfected with pMXs-IRES-blasticidin retroviral vectors with Opti-MEM and Xtreme XP Gene transfection reagent per the manufacturer’s protocol and incubated at 37°C for 24 hours. Phoenix AMPHO cells already contain the viral genes necessary for making a virus, therefore only the plasmid with the gene of interest needs to be transfected. Media containing retroviral particles was collected and stored at 4°C. Media was refreshed for a second 24-hour incubation, and then collected and pooled with the first viral media collection. Viral media was cleared of cell debris by centrifugation at 700 x g for 5 minutes followed by passage through a 0.45 micron filter (Avantor PES 25 mm 0.45 μm). Virus was aliquoted and stored at −80°C prior to transduction.

**Table 2.**
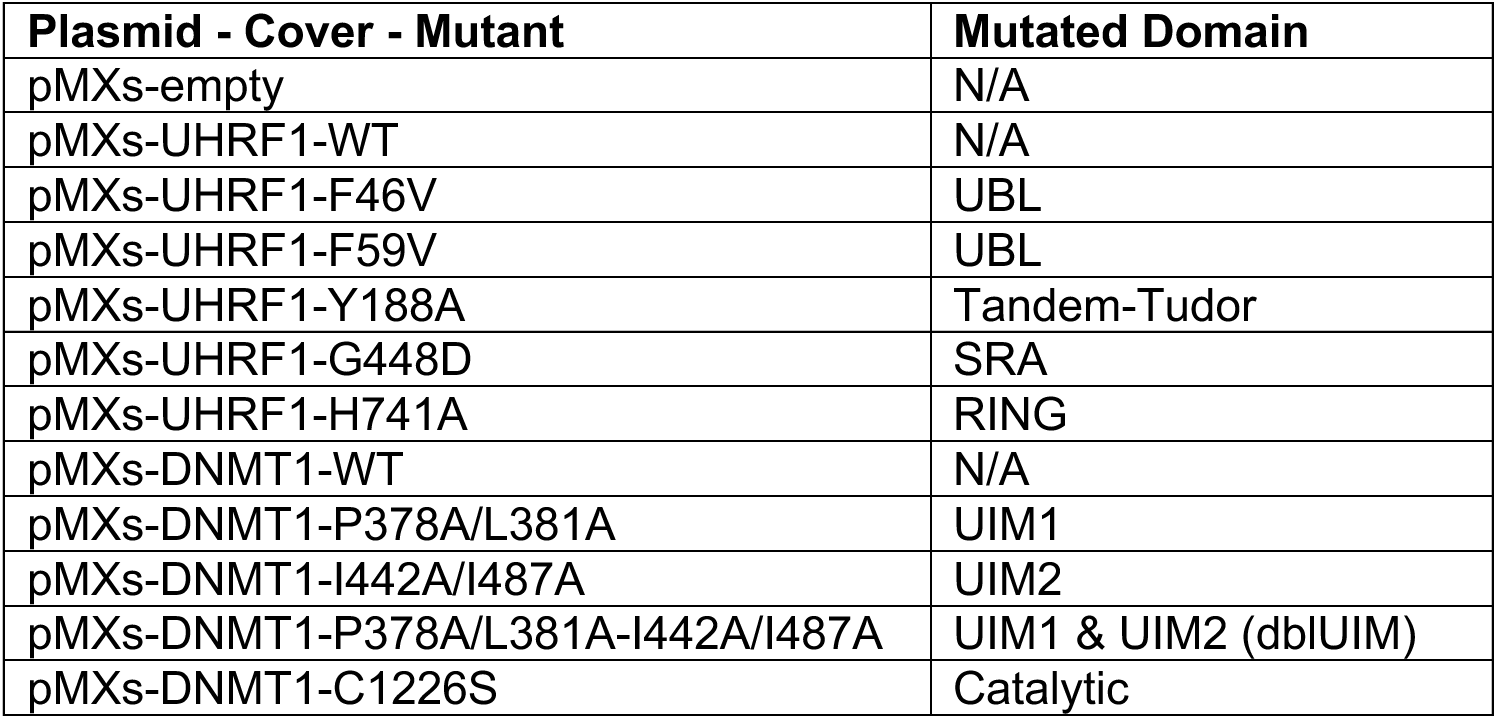
Plasmid constructs for UHRF1/DNMT1 transgene expression.

#### Transduction of HCT116 dox-inducible shUHRF1 and shDNMT1 cells

The respective shUHRF1 and shDNMT1 HCT116 cells were plated in 6-well plates and infected with media containing 8 μg/mL polybrene (with no antibiotics) and 1 ml of transgene cover retrovirus. Media was refreshed 24 hours post-infection, and the cells were allowed to grow for an additional 24 hours prior to blasticidin selection (8 μg/mL). An uninfected plate of HCT116 cells was grown in parallel to test blasticidin resistance for the infected cells, and the infected cells were maintained under blasticidin to maintain transgene cover expression.

### HCT116 doxycycline-inducible shRNA time-course

#### Doxycycline treatment

Doxycycline (Cayman) was dissolved in DMSO to a final concentration of 5 mg/mL, aliquoted, and stored at −20°C in complete darkness (dox is light sensitive). For each two-week time-course, an aliquot of dox was removed from the −20°C, thawed at room temperature, and used for the duration of the experiment. In between treatments, Dox (5 mg/mL) was stored at 4°C in complete darkness. To treat cells, Dox (5 mg/mL) was diluted in PBS (Gibco) to a final concentration of 10 ng/μl and then added to the cell culture media at a final concentration of 10 ng/mL. For the NoDox control, an equivalent amount of DMSO was applied. Dox treatments were refreshed every two days throughout the duration of the knockdown time-course.

#### Time-course

For all dox-inducible time-course experiments, the respective HCT116 shRNA cell line was passaged into two separate 10 cm cell culture plates for parallel growth in the absence (Baseline) and presence (Knockdown) of Dox (10 ng/mL). Following 2 weeks of treatment, dox was washed out and no longer added to the media. The parallel plates were maintained for an additional two (Recovery) to six weeks (for HCT116 shUHRF1 Cl.3 and Cl.6) with collection of genomic DNA and protein lysates throughout. HCT116 shGFP cell line for determining dox associated effects was a gift from Stuart Aaronson^71^.

### Western Blotting

Cells lysates were prepared as previously described^72^. Briefly, cells were lysed in CSK buffer (10 mM PIPES pH 7.0, 300 mM sucrose, 100 mM NaCl, 3 mM MgCl_2_, 0.1% Triton X-100, universal nuclease, and protease inhibitor cocktail (Roche cOmplete Mini tablets, EDTA-free)) on ice and centrifuged to remove insoluble components. Lysates were quantified and normalized using the Bradford assay (Bio-Rad), and total protein was size-separated by SDS-PAGE before semi-dry Western blot transfer to PVDF membrane. Membranes were incubated with primary antibodies against β-Actin (1:1,000; Cell Signaling Technology 4970; RRID:AB_2223172), Beta-tubulin (1:50,000; Proteintech 66240-1-Ig), DNMT1 (1:1000; Abcam ab134148) and UHRF1 (1:1,000; Cell Signaling Technology 12387; RRID:AB_2715501) for one hour. Following 3 washes in PBS-T, membranes were incubated with HRP-conjugated secondary antibody (1:10,000; Sigma-Aldrich GENA934, RRID:AB_2722659) for 1 hour at room temp. Membranes were washed again in PBS-T, incubated with ECL substrate, and imaged with film.

### DNA isolation

Cells were lysed overnight at 37°C in 2 mL of TE-SDS buffer (10 mM Tris-HCl pH 8.0, 0.1 mM EDTA, 0.5% SDS), supplemented with 100 μl of 20 mg/mL proteinase K. DNA was purified by phenol:chloroform extraction in three phases: (1) 100% phenol, (2) phenol:chloroform:isoamyl alcohol (25:24:1), and (3) chloroform:isoamyl alcohol (24:1). For each phase, the aqueous layer was combined with the organic layer in a 1:1 ratio. Samples were quickly shaken, allowed to sit on ice for approximately 5 minutes, and then separated by centrifugation at 1,693 x g for 5 minutes at 4°C. The top aqueous layer was then transferred to a new tube for the next organic phase. Following extraction, DNA was precipitated with 1/10 volume 3 M sodium acetate pH 4.8 and 2.5 volumes 100% ethanol and stored overnight at −20°C. Precipitated DNA was pelleted by centrifugation at 17,090 x g for 30 minutes at 4°C. The pelleted DNA was washed twice with 70% ethanol, allowed to dry for 15 minutes, and resuspended in TE buffer (10 mM Tris-HCl pH 8.0, 0.1 mM EDTA). Samples were then treated with 1 mg/mL RNAse A at 37°C for 30 minutes and then re-purified by ethanol precipitation as described.

### High Resolution Melt (HRM) analysis for DNA methylation

The EZ DNA Methylation Kit (Zymo D5002) was used to bisulfite convert 500 ng of DNA per sample according to the manufacturer’s protocol. Bisulfite converted DNA was eluted in 10 µl of M-elution buffer from the kit and brought up to 54 µL total with DNase-Free water. 5 µL of the bisulfite converted DNA was combined with 10 µl of Precision Melt Supermix for High Resolution Melt Analysis (BioRad 1725112) and 2 µl of Forward and Reverse primers (2 µM stock) (**Table 3**) and brought to 20 µl with DNase-Free water. A BioRAD CFX Opus93 Real-Time PCR System was used to amplify the DNA at a 60°C annealing temp for 39 cycles and then perform a melt analysis from 65 to 95°C with 0.1°C /10 sec increments. The melt temp (Tm) at the maximum reported RFU value was reported for each amplicon.

**Table 3.**
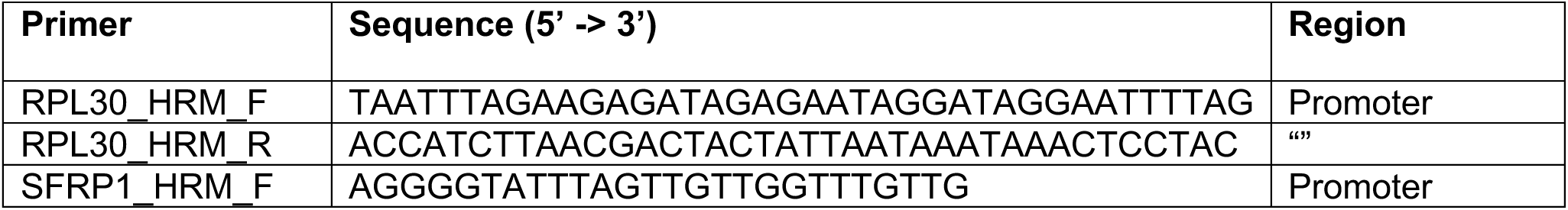
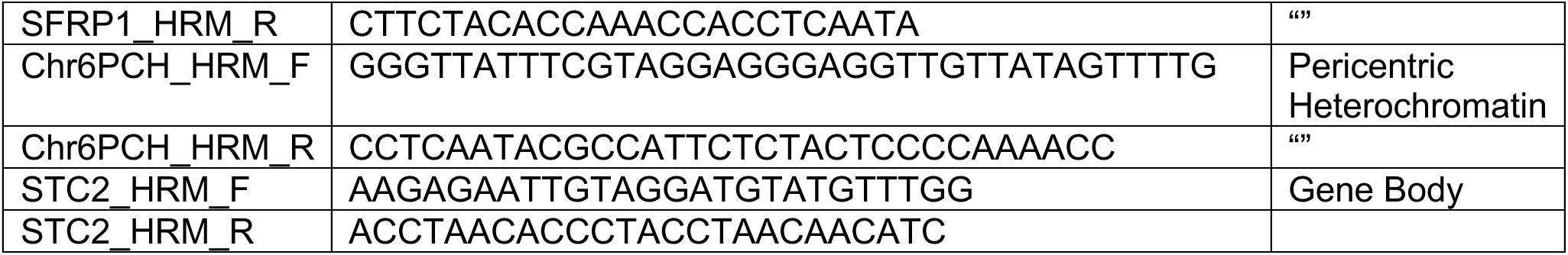
High Resolution Melt (HRM) primer sequences.

An amplicon from an unmethylated gene, *RPL30*, was used to control for bisulfite conversion. Genomic DNA isolated from HCT116 DKO1 cells (PMID: 11932749) was used as a control for determining relative degree of hypomethylation.

### Infinium MethylationEPIC BeadChip (EPIC array)

Genomic DNA was quantified by High Sensitivity Qubit Fluorometric Quantification (Invitrogen), and 1.5 μg of genomic DNA was submitted to the Van Andel Institute Genomics Core for quality control analysis, bisulfite conversion, and DNA methylation quantification using the Infinium MethylationEPIC BeadChIP (Illumina) processed on an Illumina iScan system following the manufacturer’s standard protocol (**PMID: 22122642**, **PMID: 21839163**).

### EPIC array data processing

All analyses were conducted in the R statistical software (v4.0.4) (**R Core Team**). Raw IDAT files for each sample were processed using the Bioconductor package “SeSAMe” (v1.8.12) for extraction of probe signal intensity values, normalization of probe signal intensity values, and calculation of β-values from the normalized probe signal intensity values^73–75^. The β-value is the measure of DNA methylation for each individual CpG probe, where a minimum value of 0 indicates a fully unmethylated CpG and a maximum value of 1 indicates a fully methylated CpG in the population. CpG probes with a detection p-value > 0.05 in any one sample were excluded from analyses.

### Whole Genome Bisulfite Sequencing (WGBS) sequencing and processing

200 ng of DNA was bisfulfite converted using the EZ DNA Methylation-Gold Kit (Zymo, #D5005) according to manufacturer’s instructions. Input DNA was spiked with unmethylated lambda DNA (0.5%) (Promega, #D1521). Replicate bisulfite libraries were generated with the CEGX TrueMethyl® Whole Genome Kit (CEGX, #CEGXTMWG, v3.1) according to manufacturer’s instructions. Libraries were sequenced on the Illumina X Ten. Sequencing reads from WGBS data were aligned to the human genome using v1.2 of an internally developed pipeline Meth10X^41^. This is publicly available and can be downloaded from https://github.com/luuloi/Meth10X. Total methylation levels (mC) were calculated by dividing the sum of all C calls with the sum of all C+T calls, and CpGs with a minimum coverage of 5 were used for downstream analyses. Methylation domains (PMD, LMR, UMR) were called using MethylSeekR^76^ (v1.0). HMD domains were called using the ‘complement’ function in bedtools (v2.25.0) against all the coordinates of PMDs, LMRs, and UMRs and were included if the average DNA methylation (average mC) was ≥ 0.8 across the domain.

### Determining differentially hypomethylated CpGs

Differential methylation was calculated within parallel plated experiments (for both bulk and clonal populations) as either Δβ-value or ΔmC for EPIC array and WGBS, respectively, where:

*EPIC array*

Δβ-value_Knockdown_ = β-value_Knockdown_ – β-value_Baseline_

Δβ-value_Recovery_ = β-value_Recovery_ – β-value_Baseline_

*WGBS*

ΔmC-value_Knockdown_ = mC_Knockdown_ – mC_Baseline_

ΔmC-value_Recovery_ = mC_Recovery_ – mC_Baseline_

Unless noted, we considered Δβ-value/ΔmC ≤ −0.3 as differentially hypomethylated as this is interpreted as roughly 30% reduction in DNA methylation relative to the baseline CpG methylation value.

### Enrichment Bias Calculation and Hypergeometic Distribution Testing

CpGs (both EPIC and WGBS data) were mapped to their genomic coordinate (hg38) and were then annotated to their genomic annotation relationship (e.g. promoter-TSS, exon) using HOMER^77^ (v4.10.3). All enrichment bias calculations were normalized to the distribution of all highly methylated CpGs (within individual bulk and clonal cell populations) on the EPIC array (β-value_Baseline_ ≥ 0.85) and WGBS (mC_Baseline_ ≥ 0.85), respectively.

Enrichment bias calculations were done by first determining the following values for each feature:

*q* = Number of CpGs that are differentially methylated in feature (e.g. low-density CpGs)

*m* = Total number of highly methylated CpGs that match feature (e.g. all highly methylated low-density CpGs)

*n* = Total number CpGs that do not match feature (e.g. all highly methylated high-density CpGs)

*k* = Total number of all differentially methylated CpGs (e.g. low-density CpGs + high-density CpGs)

Next, the expected number of CpGs that would be differentially methylated in that feature by random chance was determined with the following equation:

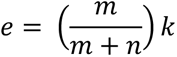

Finally, percent enrichment bias was calculated with the following equation:

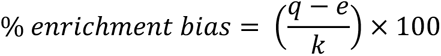

Where positive or negative enrichment values indicate more or less enrichment for a feature than would be expected by random chance, respectively.

Hypergeometric distribution testing for determining significance of enrichment bias was performed using the phyper() function in R with the following values: *q,m,n,k*.

### Repli-seq Integration

16-phase RepliSeq data (measuring replication timing from early to late replication) for HCT116 was downloaded^78^ (GEO: GSE137764) and each phase of replication timing was separated into their own genomic coordinates (both bed and bigwig files) for use in integrative DNA methylation analysis.

### Methylation array analysis of publicly available data

Raw idat files were downloaded from the Gene Expression Omnibus under accession GSE68379 (additional cell lines) and GSE118970 (HCT116/RKO UHRF1 mutants). All analyses were conducted in the R statistical software (v4.0.4) as described under **EPIC array data processing.**

### Chromatin Immunoprecipitation

#### Cell Fixation and Collection

Approximately 10 million HCT116 shUHRF1 Clone 3 and Clone 6 NoDox cells were grown to 80% confluency in a 10 cm plates. Cells were washed in the plate with 5 mL of 1X PBS at room temperature. The 1X PBS wash was removed, and 5 mL of Fixing Buffer (50 mM HEPES-KOH pH 7.6, 100 mM NaCl, 1 mM EDTA pH 8.0, 0.5 mM EGTA pH 8.0) was added. Cells were fixed by adding 313 μl of freshly prepared 16% methanol-free formaldehyde solution (Thermo Scientific Catalog #28906) to a final concentration of ∼1%, and cells were incubated on a shaker at room temperature for 10 minutes. Formaldehyde was quenched by adding 266 μl of 2.5 M Glycine (final concentration of 125 mM), and cells were incubated for an additional 5 minutes on the shaker at room temperature. Cells were collected from the plate by scraping the monolayer and diluting the fixation solution with 10 mL ice-cold 1X PBS. Cells were pelleted at 200 x g for 5 minutes at 4°C. Cells were washed twice more with 5 mL ice-cold 1X PBS and collected by centrifugation at 200 x g for 5 minutes at 4°C. The final PBS wash was carefully aspirated from the cell pellet, and cells were flash frozen in liquid nitrogen and stored at −80°C until use.

#### Nuclei isolation and sonication

Cells were thawed on ice for 10 minutes prior to cell lysis. Cells were lysed in 1 mL LB1 (50 mM HEPES-KOH pH 7.6, 140 mM NaCl, 1 mM EDTA, 10% Glycerol, 0.5% NP-40, 0.25% Triton X-100, Protease inhibitor cocktail (Roche cOmplete Mini tablets, EDTA-free)) and incubated for 10 minutes rotating at 4°C. Intact nuclei were collected by centrifugation at 1,700 x g for 5 minutes at 4°C. Supernatant was removed so as not to disturb the nuclei pellet, and nuclei were resuspended and washed in 1 mL LB2 (10 mM Tris-HCl pH 8.0, 1 mM EDTA, 0.5 mM EGTA, 200 mM NaCl, protease cocktail inhibitor) for 10 minutes rotating at 4°C. Nuclei were collected by centrifugation at 1,700 x g for 5 minutes at 4°C, and the supernatant was removed so as not to disturb the nuclei pellet. Finally, nuclei pellets were gently rinsed twice without disturbing the pellet with 1 mL LB3 (10 mM Tris-HCl pH 8.0, 1 mM EDTA, 0.5 mM EGTA, 0.01% NP-40, protease cocktail inhibitor) and collected at 1,700 x g for 5 minutes at 4°C. Following the two rinse steps, nuclei were resuspended in 1 mL LB3 and transferred to a 1 mL milliTUBE (Covaris) for shearing. Nuclei were lysed and chromatin was sheared to range of 300-600 base-pair fragments using a Covaris E220 evolution Focused ultrasonicator with the following parameters: Peak power (140.0), Duty Factor (5.0), Cycles/Burst (200), Duration (600 seconds), Temperature (4°C).

#### Immunoprecipitation

Sheared chromatin was quantified by Bradford Assay, and 300 μg of chromatin was brought to a final volume of 500 μl in LB3, and then an additional 500 μl of ChIP Cocktail Mix (40 mM Tris-HCl pH 7.6, 150 mM NaCl, 1 mM EDTA pH 8.0, 1% Triton X-100, 0.5% NP-40, Protease inhibitor cocktail) was added to bring the final volume to 1 mL. Prepared chromatin was then pre-cleared by incubation with 20 μl of pre-washed Dynabeads Protein G magnetic beads (Invitrogen Catalog #:10004D) for 3 hours at 4°C with constant rotation. Prior to incubation with antibody, 10 μl of pre-cleared chromatin was removed and set aside to serve as 1% input. Pre-cleared chromatin was removed by magnetic separation, transferred to a new tube, and immunoprecipitated with either 5 μl of H3K9me3 antibody (Active Motif 39161, Lot# 14418003) or 5 μl of H3K27me3 antibody (Cell Signaling C36B11, Lot#:97335(14)) overnight at 4°C with constant rotation. Protein G magnetic beads (35 μl/IP) were blocked overnight in 1 mL of 1X PBS, 0.5% BSA, and 20 μg of Herring Sperm DNA (Sigma Catalog #D7290) at 4°C with constant rotation. The next morning, blocked beads were washed three times with 1X PBS + 0.5% BSA, and two times with WB1 (50 mM Tris-HCl pH 7.6, 150 mM NaCl, 5 mM EDTA pH 8.0, 0.5% NP-40, 1% Triton X-100). Antibody-chromatin complexes were incubated with blocked beads for 3 hours at 4°C with constant rotation. Unbound chromatin was then removed using magnetic separation, and the beads were washed as follows: 3 times with WB1, 3 times with WB2 (50 mM Tris-HCl pH 7.6, 500 mM NaCl, 5 mM EDTA pH 8.0, 0.5% NP-40, 1% Triton X-100), 2 times with WB1, and 1 time with Low Salt TE (10 mM Tris-HCl pH 8.0, 1 mM EDTA pH 8.0, 50 mM NaCl). Each wash was a 5 minute incubation at 4°C with constant rotation followed by magnetic separation and removal of buffer.

#### Elution and DNA clean-up

To elute DNA from the magnetic beads, 50 μl of Elution Buffer (10 mM Tris-HCl pH 8.0, 10 mM EDTA, 150 mM NaCl, 5 mM DTT, 1% SDS) was added to the beads and incubated at 65°C for 15 minutes. The elution step was repeated, and eluates combined. Eluents and 1% input (with 90 μl of elution buffer added) were incubated overnight at 65°C with constant shaking to reverse crosslink protein:DNA complexes. The next morning, 2 μl of DNase-free RNase A (10 mg/mL stock) was added to eluents and incubated at 37°C for 1 hour. Next, 10 μl of Proteinase K (20 mg/mL stock) was added to eluents and incubated at 37°C for 2 hours. DNA was isolated following standard KAPA Pure Beads (KAPA Biosystems Catalog# KK8000) protocol with a 1.5X ratio of beads to DNA. Final elution was in 20 μl of Nuclease-free water, and DNA concentration was measure by Qubit dsDNA High Sensitivity Assay kit (ThermoFisher Scientific Catalog#: Q32851).

#### Library preparation

Immunoprecipitated fragments and saved inputs were quantified by High Sensitivity Qubit Fluorometric Quantification (Invitrogen), and 10 ng of purified DNA for each IP and input sample were used for library preparation with the KAPA Hyper Prep Kit (Part#: KR0961) for these samples.

Library preparation including fragment end-repair, A-tail extension, and adapter ligation was conducted per the manufacturer’s instructions (KAPA). Adapter-ligated fragments were amplified with 11 cycles following the recommended thermocycler program, and DNA was purified with two rounds of purification using KAPA Pure Beads (#KK8000). Quality and quantity of the finished libraries were assessed using a combination of Agilent DNA High Sensitivity chip (Agilent Technologies, Inc.), QuantiFluor® dsDNA System (Promega Corp., Madison, WI, USA), and Kapa Illumina Library Quantification qPCR assays (Kapa Biosystems). Individually indexed libraries were pooled and 50 bp, paired end sequencing was performed on an Illumina NovaSeq6000 sequencer using an S2, 100 bp sequencing kit to a minimum read depth of 50M read pairs per IP library and 100M read pairs per Input library. Base calling was done by Illumina RTA3 and output of NCS was demultiplexed and converted to FastQ format with Illumina Bcl2fastq (v1.9.0).

### ChIP-seq processing and analysis

ChIP sequencing reads were 3’ trimmed and filtered for quality and adapter content using TrimGalore (v0.5.0) and quality was assessed by FastQC (v0.11.8). Reads were aligned to human genome assembly hg38 with bowtie2 (v2.3.5) and were deduplicated using removeDups from samblaster^79^ (v.0.1.24). Aligned BAM files were used for quality control analysis with deeptools^80^ (v3.2.0) ‘plotFingerprint’ and ‘plotPCA’ functions. As both H3K9me3 and H3K27me3 domains are broad, we called peaks using Enriched Domain Detector (EDD) with default parameters^81^. For HCT116 shUHRF1 Clone 3 and Clone 6 cells (NoDox, baseline), H3K9me3 and H3K27me3 peak coordinates that were consistent between biological replicates were used for downstream analysis. For shUHRF1 Clone 9 cells and shDNMT1 bulk cell populations, peak coordinates that were consistent between both Clone 3 and Clone 6 (NoDox, baseline) cells were used. For HCT116 H3K36me3 distributions, we used publicly available data (ENCSR091QXP, GSE95914).

### Integrative genomic analysis

To integrate WGBS DNA methylation data with genomic annotations of interest (Replication timing (Repli-seq), H3K9me3/H3K27me3/H3K36me3 distributions (ChIP-seq)), bigwig files for mC-values and ΔmC-values (WGBS) were generated for each sample and time-point for all CpGs, high-density CpGS, and low-density CpGs. Bed files with genomic coordinates for H3K9me3, H3K27me3, and Repli-seq phases were generated as described above. Integrated analyses were conducted with deeptools^80^ (v3.2.0) by constructing matrices with ‘computeMatrix’ across queried genomic coordinates with the respective bigwig data and calculating the average mC-values and ΔmC-values with ‘plotProfile’ and ‘plotHeatmap’.

### Methyl Domain distribution overlap analysis

Genomic coordinates for Methyl Domains across samples and time-points were determined as described in **Whole Genome Bisulfite Sequencing (WGBS) sequencing and processing.** Genomic coverage of HMDs, PMDs, LMRs, and UMRs were calculated by summation of the length (in base pairs) for each respective domain within each individual sample. To determine transition coverage, Baseline HMD and PMD genomic coordinates were intersected with bedtools^82^ (v2.25.0) ‘intersect’ command with the genomic coordinates for Methyl Domains in the Knockdown and Recovery samples, and the length of the intersected domains were summed. Additional overlap analyses among the Methyl Domain genomic locations were conducted using the ‘jaccard’ command from bedtools.

### Protein purification

Recombinant UHRF1 and DNMT1 were generated and purified as full-length proteins with N-terminal 6x-His-MBP (maltose binding protein) tags as previously described^46^. DNMT1 RFTS domain wild type and mutants were cloned into a pQE-80L plasmid with an N-terminal 6x-His-MBP tag. Plasmids were transformed into BL21 DE3 *E. coli* and grown at 37°C to an OD_600_ of 1.0, followed by addition of 0.2 mM IPTG and induction overnight at 16°C. Induced cultures were harvested and bacteria were resuspended in lysis buffer (50 mM HEPES pH 7.5, 250 mM NaCl, 20 mM imidazole, 30 uM ZnOAc, 1 mM PMSF, 1 mM DTT). Cells were lysed by addition of lysozyme and sonication. Lysate was cleared by centrifugation at 38,000 x g at 4°C for 30 minutes. Lysates were incubated with His60 Ni Superflow resin (Takara Bio) at 4°C with rotation for 1 hour. Resin was washed 3x with at least 10 volumes of lysis buffer followed by elution of bound proteins by addition of 5 volumes of elution buffer (25 mM HEPES pH 7.5, 100 mM NaCl, 250 mM imidazole, 1 mM DTT). Protein purity was assessed by SDS-PAGE and Coomassie staining. Purified proteins were dialyzed into storage buffer (25mM HEPES pH 7.5, 100mM NaCl, 10% glycerol, 1 mM DTT), concentrated via centrifugal filtration, snap frozen in small aliquots and stored at −80°C.

### Fluorescence Polarization

Fluorescence polarization binding assays were performed as described^83^. Briefly, recombinant UHRF1 was titrated into assay buffer (25 mM HEPES pH 7.5, 100 mM NaCl, 0.05% NP-40) containing 10 nM fluorescent DNA oligonucleotides (5’-FAM-CCAT**X**(5mC)G**X**TGAC-3’ where **X** was interchanged with specific flanking nucleotides (**G**CG**C**, **C**GC**C**, **C**CG**G**; **A**CG**T**, **A**CG**A**, **T**CG**A**)). Measurements were performed in 384-well plates with triplicate 25 µL reactions and plotted as change in anisotropy.

### Construction of H3K23ub mononucleosomes

H3 “K23ub” thioether mimic was synthesized following a method similar to that described by Hann *et. al*^84^. Briefly, a maleimide moiety was covalently conjugated to the C-terminus of ubiquitin (Ub-mal). This Ub-mal was then conjugated to H3 K23C through a stable thioether linkage. The H3 “K23ub” thioether mimic was then assembled into histone octamers and subsequently mononucleosomes. The syntheses of intermediates and final products are described below.

#### Synthesis of ubiquitin-thioester (UbTE)

Ubiquitin-GyraseA-His_6_ (UbGyrA) was expressed in Rosetta *E. coli* (RDE3) and induced at 16 °C overnight. The pellet was resuspended in lysis buffer (50 mM potassium phosphate monobasic, 300 mM NaCl, 20 mM imidazole, 1 mM PMSF, pH 8.0) and lysed using a rod sonicator. The lysate was cleared by centrifugation and the supernatant was loaded onto Ni-NTA beads. The beads were then washed three time with wash buffer (50 mM potassium phosphate monobasic, 1 M NaCl, 20 mM imidazole, 1 % NP-40, pH 8.0) two times with lysis buffer, and eluted in 1x PBS, 500 mM imidazole, 1 mM TCEP, pH 7. Solid MESNa was added to the eluate to a final concentration of 500 mM and the pH was readjusted to 7. The reaction was nutated at 4 °C for 48 hours and splicing was monitored by LCMS. The reaction was then dialyzed against 1 % AcOH, the precipitate was removed by centrifugation, and the supernatant was lyophilized. The crude ubiquitin-thioester (UbTE) was purified by reversed-phase HPLC on a 0-70 % buffer B gradient (HPLC solvent A = 0.1 % TFA in ddH_2_O, HPLC solvent B = 90 % ACN, 10 % ddH_2_O, 0.1 % TFA). Pooled fractions were lyophilized and stored at −80 °C.

#### Synthesis of ubiquitin-acyl hydrazide (Ub-NHNH_2_)

UbTE was dissolved in 1x PBS, 6 M guanidine, pH 5, and hydrazine was added to a final concentration of 500 mM. The reaction was incubated at 30 °C for 30 minutes and then immediately diluted five-fold into HPLC solvent A. The Ub-NHNH_2_ was then purified by reversed-phase HPLC on a 0-70 % solvent B gradient. Pooled fractions were lyophilized and stored at −80 °C.

#### Synthesis of ubiquitin-N-(2-aminoethyl)maleimide (Ub-mal)

Ub-NHNH_2_ was dissolved in oxidation buffer (0.2 M phosphate, 20 mM NaNO_3_, 6 M guanidine, pH 3) and nutated at 4 °C to form the acyl-azide *in situ*. An equal volume of 200 mM N-(2-aminoethyl)maleimide dissolved in 1.5 M phosphate pH 7.4 was added to the acyl-azide and the reaction was nutated at room temperature for 15 minutes to form the Ub-mal. The reaction was diluted five-fold into HPLC buffer A and Ub-mal was purified by reversed-phase HPLC on a 0-70 % solvent B gradient. Pooled fractions were lyophilized and stored at −80 °C.

#### Synthesis of H3 K23ub thioether mimic

An excess of H3 K23C was combined with Ub-mal and dissolved in 1x PBS, pH 6.8. The reaction was nutated at 4 °C overnight and conversion was monitored by LCMS. Upon complete consumption of free Ub-mal, the reaction was diluted five-fold into HPLC solvent A and H3K23ub purified by reversed-phase HPLC on a 30-70 % solvent B gradient. Pooled fractions were lyophilized and stored at −80 °C.

#### Histone octamer assembly

Histones were dissolved in unfolding buffer (20 mM tris, 6 M guanidine, 0.5 mM EDTA, 10 mM TCEP, pH 7.5 and combined in a 1:1:0.95:0.95 molar ratio (H2A:H2B:H3K23ub:H4). The pooled histone solution was diluted with unfolding buffer to a final combined histone concentration of 1 mg/mL. The histone mixture was then dialyzed against refolding buffer (10 mM tris, 2 M NaCl, 0.5 mM EDTA, 1 mM TCEP, pH 7.5) multiple times to ensure complete buffer exchange. The refolded histone octamers were concentrated and then isolated by purification on a GE Superdex 75 15/300 increase size-exclusion column. Pooled fractions were concentrated, diluted to 50 % glycerol, and stored at −20 °C.

#### Mononucleosome assembly

Mononucleosome assembly was performed according to the previously described salt dilution method with slight modification. Briefly, purified octamers were mixed with Widom-601 DNA (1:1 ratio) in a 2 M salt solution (10 mM Tris pH 7.5, 2 M NaCl, 1 mM EDTA, 1 mM DTT). After incubation at 37 °C for 15 min, the mixture was gradually diluted (∼15 min) at 30 °C with dilution buffer (10 mM Tris pH 7.5, 10 mM NaCl, 1 mM EDTA, 1 mM DTT). The assembled mononucleosomes were concentrated and characterized by native gel electrophoresis (5% acrylamide gel, 0.5x TBE, 120 V, 40 min) using ethidium bromide staining.

### *In vitro* methyltransferase assays

*In vitro* methyltransferase assays were performed by reacting wild type and mutant forms of recombinant His-MBP-DNMT1 with ^3^H-labeled S-adenosylmethionine and the previously described biotinylated hemi-methylated Dup_1 DNA substrate (https://doi.org/10.1093/nar/gkt753) (5’-GATmCGCmCGATGmCGmCGAATmCGmCGATmCGATGmCGAT-3’). Briefly, 10 μL reactions were assembled in DNMT1 reaction buffer (20 mM HEPES pH 7.5, 50 mM KCl, 1 mM EDTA, 25 μg/mL BSA) containing 150 nM DNMT1, 300 nM DNA, and 20 μM ^3^H-SAM. Reactions were incubated at 37°C, then quenched at various timepoints by addition of 190 μL of 20 μM S-adenosylhomocysteine (SAH).

Reactions were transferred to 96-well Streptavidin FlashPlates (Perkin Elmer) and incubated at room temp for 15 minutes to bind biotinylated DNA, followed by 3 washes with 200 μL wash buffer per well (50 mM Tris pH 7.4, 0.05% Tween-20). Scintillation counts were measured using a MicroBeta2 microplate reader (Perkin Elmer). Data was plotted using GraphPad Prism.

### Alpha screen assays

Alpha screen assays were performed in 384-well plates using the Alpha Screen Histidine (Nickel Chelate) detection kit from Revvity. H3K14ub and H3K18ub nucleosomes (with native ubiquitin linkages) were purchased from Epicypher. H3K23ub nucleosomes (with DCA mimic ubiquitin linkage) were synthesized as described. Wild-type and mutant recombinant His-MBP-RFTS as well as ubiquitinated nucleosomes (with 5’ biotinylated DNA) were diluted in alpha assay buffer (25mM HEPES pH 7.5, 250mM NaCl, 0.05% NP40), mixed in a final volume of 10 μL, and incubated at room temperature for 30 minutes. Alpha donor and acceptor beads were diluted in alpha assay buffer and added to the reactions. The final reaction volume was 20 μL and the final concentration of each bead was 20 μg/mL. Reactions were incubated at room temperature for 30 minutes protected from light.

Plates were read on an alpha screen compatible Perkin Elmer plate reader. Data was plotted in GraphPad Prism and fitted with nonlinear regression using the specific binding with Hill slope mod.

### Materials availability

All unique/stable reagents (plasmids, cell lines) generated in this study are available from the Lead Contact with a completed Materials Transfer Agreement.

### Data and code availability

All WGBS, ChIP-seq, and EPIC array raw and processed data will be deposited in the Gene Expression Omnibus (GEO) archive and will be released to the public upon publication. All code used for processing, analysis, and figure generation will be deposited on our GitHub site (https://github.com/rleetied) and archived on Zenodo prior to publication.

## REFERENCES

1. Bird, A.P. (1980). DNA methylation and the frequency of CpG in animal DNA. Nucleic Acids Res. 8, 1499–1504. 10.1093/nar/8.7.1499.

2. Colaneri, A., Staffa, N., Fargo, D.C., Gao, Y., Wang, T., Peddada, S.D., and Birnbaumer, L. (2011). Expanded methyl-sensitive cut counting reveals hypomethylation as an epigenetic state that highlights functional sequences of the genome. Proc. Natl. Acad. Sci. 108, 9715–9720. 10.1073/pnas.1105713108.

3. Lister, R., Pelizzola, M., Dowen, R.H., Hawkins, R.D., Hon, G., Tonti-Filippini, J., Nery, J.R., Lee, L., Ye, Z., Ngo, Q.-M., et al. (2009). Human DNA methylomes at base resolution show widespread epigenomic differences. Nature 462, 315–322. 10.1038/nature08514.

4. Ziller, M.J., Gu, H., Müller, F., Donaghey, J., Tsai, L.T.-Y., Kohlbacher, O., De Jager, P.L., Rosen, E.D., Bennett, D.A., Bernstein, B.E., et al. (2013). Charting a dynamic DNA methylation landscape of the human genome. Nature 500, 477–481. 10.1038/nature12433.

5. Baylin, S.B., and Jones, P.A. (2016). Epigenetic Determinants of Cancer. Cold Spring Harb. Perspect. Biol. 8, a019505. 10.1101/cshperspect.a019505.

6. Hansen, K.D., Timp, W., Bravo, H.C., Sabunciyan, S., Langmead, B., McDonald, O.G., Wen, B., Wu, H., Liu, Y., Diep, D., et al. (2011). Increased methylation variation in epigenetic domains across cancer types. Nat. Genet. 43, 768–775. 10.1038/ng.865.

7. Berman, B.P., Weisenberger, D.J., Aman, J.F., Hinoue, T., Ramjan, Z., Liu, Y., Noushmehr, H., Lange, C.P.E., van Dijk, C.M., Tollenaar, R.A.E.M., et al. (2012). Regions of focal DNA hypermethylation and long-range hypomethylation in colorectal cancer coincide with nuclear lamina-associated domains. Nat. Genet. 44, 40–46. 10.1038/ng.969.

8. Johnstone, S.E., Reyes, A., Qi, Y., Adriaens, C., Hegazi, E., Pelka, K., Chen, J.H., Zou, L.S., Drier, Y., Hecht, V., et al. (2020). Large-Scale Topological Changes Restrain Malignant Progression in Colorectal Cancer. Cell 182, 1474–1489.e23. 10.1016/j.cell.2020.07.030.

9. Salhab, A., Nordström, K., Gasparoni, G., Kattler, K., Ebert, P., Ramirez, F., Arrigoni, L., Müller, F., Polansky, J.K., Cadenas, C., et al. (2018). A comprehensive analysis of 195 DNA methylomes reveals shared and cell-specific features of partially methylated domains. Genome Biol. 19, 150. 10.1186/s13059-018-1510-5.

10. Brinkman, A.B., Nik-Zainal, S., Simmer, F., Rodríguez-González, F.G., Smid, M., Alexandrov, L.B., Butler, A., Martin, S., Davies, H., Glodzik, D., et al. (2019). Partially methylated domains are hypervariable in breast cancer and fuel widespread CpG island hypermethylation. Nat. Commun. 10, 1749. 10.1038/s41467-019-09828-0.

11. Zhou, W., Dinh, H.Q., Ramjan, Z., Weisenberger, D.J., Nicolet, C.M., Shen, H., Laird, P.W., and Berman, B.P. (2018). DNA methylation loss in late-replicating domains is linked to mitotic cell division. Nat. Genet. 50, 591–602. 10.1038/s41588-018-0073-4.

12. Gaidatzis, D., Burger, L., Murr, R., Lerch, A., Dessus-Babus, S., Schübeler, D., and Stadler, M.B. (2014). DNA Sequence Explains Seemingly Disordered Methylation Levels in Partially Methylated Domains of Mammalian Genomes. PLOS Genet. 10, e1004143. 10.1371/journal.pgen.1004143.

13. Decato, B.E., Qu, J., Ji, X., Wagenblast, E., Knott, S.R.V., Hannon, G.J., and Smith, A.D. (2020). Characterization of universal features of partially methylated domains across tissues and species. Epigenetics Chromatin 13, 39. 10.1186/s13072-020-00363-7.

14. Hon, G.C., Hawkins, R.D., Caballero, O.L., Lo, C., Lister, R., Pelizzola, M., Valsesia, A., Ye, Z., Kuan, S., Edsall, L.E., et al. (2012). Global DNA hypomethylation coupled to repressive chromatin domain formation and gene silencing in breast cancer. Genome Res. 22, 246–258. 10.1101/gr.125872.111.

15. Endicott, J.L., Nolte, P.A., Shen, H., and Laird, P.W. (2022). Cell division drives DNA methylation loss in late-replicating domains in primary human cells. Nat. Commun. 13, 6659. 10.1038/s41467-022-34268-8.

16. Du, Q., Bert, S.A., Armstrong, N.J., Caldon, C.E., Song, J.Z., Nair, S.S., Gould, C.M., Luu, P.-L., Peters, T., Khoury, A., et al. (2019). Replication timing and epigenome remodelling are associated with the nature of chromosomal rearrangements in cancer. Nat. Commun. 10, 416. 10.1038/s41467-019-08302-1.

17. Sharif, J., Muto, M., Takebayashi, S., Suetake, I., Iwamatsu, A., Endo, T.A., Shinga, J., Mizutani-Koseki, Y., Toyoda, T., Okamura, K., et al. (2007). The SRA protein Np95 mediates epigenetic inheritance by recruiting Dnmt1 to methylated DNA. Nature 450, 908–912. 10.1038/nature06397.

18. Bostick, M., Kim, J.K., Estève, P.-O., Clark, A., Pradhan, S., and Jacobsen, S.E. (2007). UHRF1 Plays a Role in Maintaining DNA Methylation in Mammalian Cells. Science 317, 1760–1764. 10.1126/science.1147939.

19. Feng, S., Cokus, S.J., Zhang, X., Chen, P.-Y., Bostick, M., Goll, M.G., Hetzel, J., Jain, J., Strauss, S.H., Halpern, M.E., et al. (2010). Conservation and divergence of methylation patterning in plants and animals. Proc. Natl. Acad. Sci. 107, 8689–8694. 10.1073/pnas.1002720107.

20. Petryk, N., Bultmann, S., Bartke, T., and Defossez, P.-A. (2021). Staying true to yourself: mechanisms of DNA methylation maintenance in mammals. Nucleic Acids Res. 49, 3020–3032. 10.1093/nar/gkaa1154.

21. Rothbart, S.B., Krajewski, K., Nady, N., Tempel, W., Xue, S., Badeaux, A.I., Barsyte-Lovejoy, D., Martinez, J.Y., Bedford, M.T., Fuchs, S.M., et al. (2012). Association of UHRF1 with methylated H3K9 directs the maintenance of DNA methylation. Nat. Struct. Mol. Biol. 19, 1155–1160. 10.1038/nsmb.2391.

22. Ferry, L., Fournier, A., Tsusaka, T., Adelmant, G., Shimazu, T., Matano, S., Kirsh, O., Amouroux, R., Dohmae, N., Suzuki, T., et al. (2017). Methylation of DNA Ligase 1 by G9a/GLP Recruits UHRF1 to Replicating DNA and Regulates DNA Methylation. Mol. Cell 67, 550–565.e5. 10.1016/j.molcel.2017.07.012.

23. Rothbart, S.B., Dickson, B.M., Ong, M.S., Krajewski, K., Houliston, S., Kireev, D.B., Arrowsmith, C.H., and Strahl, B.D. (2013). Multivalent histone engagement by the linked tandem Tudor and PHD domains of UHRF1 is required for the epigenetic inheritance of DNA methylation. Genes Dev. 27, 1288–1298. 10.1101/gad.220467.113.

24. Han, M., Li, J., Cao, Y., Huang, Y., Li, W., Zhu, H., Zhao, Q., Han, J.-D.J., Wu, Q., Li, J., et al. (2020). A role for LSH in facilitating DNA methylation by DNMT1 through enhancing UHRF1 chromatin association. Nucleic Acids Res. 48, 12116–12134. 10.1093/nar/gkaa1003.

25. Liu, X., Gao, Q., Li, P., Zhao, Q., Zhang, J., Li, J., Koseki, H., and Wong, J. (2013). UHRF1 targets DNMT1 for DNA methylation through cooperative binding of hemi-methylated DNA and methylated H3K9. Nat. Commun. 4, 1563. 10.1038/ncomms2562.

26. Dennis, K., Fan, T., Geiman, T., Yan, Q., and Muegge, K. (2001). Lsh, a member of the SNF2 family, is required for genome-wide methylation. Genes Dev. 15, 2940–2944. 10.1101/gad.929101.

27. Arita, K., Isogai, S., Oda, T., Unoki, M., Sugita, K., Sekiyama, N., Kuwata, K., Hamamoto, R., Tochio, H., Sato, M., et al. (2012). Recognition of modification status on a histone H3 tail by linked histone reader modules of the epigenetic regulator UHRF1. Proc. Natl. Acad. Sci. U. S. A. 109, 12950–12955. 10.1073/pnas.1203701109.

28. Avvakumov, G.V., Walker, J.R., Xue, S., Li, Y., Duan, S., Bronner, C., Arrowsmith, C.H., and Dhe-Paganon, S. (2008). Structural basis for recognition of hemi-methylated DNA by the SRA domain of human UHRF1. Nature 455, 822–825. 10.1038/nature07273.

29. Arita, K., Ariyoshi, M., Tochio, H., Nakamura, Y., and Shirakawa, M. (2008). Recognition of hemi-methylated DNA by the SRA protein UHRF1 by a base-flipping mechanism. Nature 455, 818–821. 10.1038/nature07249.

30. Fang, J., Cheng, J., Wang, J., Zhang, Q., Liu, M., Gong, R., Wang, P., Zhang, X., Feng, Y., Lan, W., et al. (2016). Hemi-methylated DNA opens a closed conformation of UHRF1 to facilitate its histone recognition. Nat. Commun. 7, 11197. 10.1038/ncomms11197.

31. Harrison, J.S., Cornett, E.M., Goldfarb, D., DaRosa, P.A., Li, Z.M., Yan, F., Dickson, B.M., Guo, A.H., Cantu, D.V., Kaustov, L., et al. (2016). Hemi-methylated DNA regulates DNA methylation inheritance through allosteric activation of H3 ubiquitylation by UHRF1. eLife 5, e17101. 10.7554/eLife.17101.

32. Nishiyama, A., Yamaguchi, L., Sharif, J., Johmura, Y., Kawamura, T., Nakanishi, K., Shimamura, S., Arita, K., Kodama, T., Ishikawa, F., et al. (2013). Uhrf1-dependent H3K23 ubiquitylation couples maintenance DNA methylation and replication. Nature 502, 249–253. 10.1038/nature12488.

33. Misaki, T., Yamaguchi, L., Sun, J., Orii, M., Nishiyama, A., and Nakanishi, M. (2016). The replication foci targeting sequence (RFTS) of DNMT1 functions as a potent histone H3 binding domain regulated by autoinhibition. Biochem. Biophys. Res. Commun. 470, 741–747. 10.1016/j.bbrc.2016.01.029.

34. Ishiyama, S., Nishiyama, A., Saeki, Y., Moritsugu, K., Morimoto, D., Yamaguchi, L., Arai, N., Matsumura, R., Kawakami, T., Mishima, Y., et al. (2017). Structure of the Dnmt1 Reader Module Complexed with a Unique Two-Mono-Ubiquitin Mark on Histone H3 Reveals the Basis for DNA Methylation Maintenance. Mol. Cell 68, 350–360.e7. 10.1016/j.molcel.2017.09.037.

35. Qin, W., Wolf, P., Liu, N., Link, S., Smets, M., Mastra, F.L., Forné, I., Pichler, G., Hörl, D., Fellinger, K., et al. (2015). DNA methylation requires a DNMT1 ubiquitin interacting motif (UIM) and histone ubiquitination. Cell Res. 25, 911–929. 10.1038/cr.2015.72.

36. Hermann, A., Goyal, R., and Jeltsch, A. (2004). The Dnmt1 DNA-(cytosine-C5)-methyltransferase methylates DNA processively with high preference for hemimethylated target sites. J. Biol. Chem. 279, 48350–48359. 10.1074/jbc.M403427200.

37. Vilkaitis, G., Suetake, I., Klimašauskas, S., and Tajima, S. (2005). Processive Methylation of Hemimethylated CpG Sites by Mouse Dnmt1 DNA Methyltransferase*. J. Biol. Chem. 280, 64–72. 10.1074/jbc.M411126200.

38. Goyal, R., Reinhardt, R., and Jeltsch, A. (2006). Accuracy of DNA methylation pattern preservation by the Dnmt1 methyltransferase. Nucleic Acids Res. 34, 1182–1188. 10.1093/nar/gkl002.

39. Mishima, Y., Brueckner, L., Takahashi, S., Kawakami, T., Otani, J., Shinohara, A., Takeshita, K., Garvilles, R.G., Watanabe, M., Sakai, N., et al. (2020). Enhanced processivity of Dnmt1 by monoubiquitinated histone H3. Genes Cells 25, 22–32. 10.1111/gtc.12732.

40. Pidsley, R., Zotenko, E., Peters, T.J., Lawrence, M.G., Risbridger, G.P., Molloy, P., Van Djik, S., Muhlhausler, B., Stirzaker, C., and Clark, S.J. (2016). Critical evaluation of the Illumina MethylationEPIC BeadChip microarray for whole-genome DNA methylation profiling. Genome Biol. 17, 208. 10.1186/s13059-016-1066-1.

41. Nair, S.S., Luu, P.-L., Qu, W., Maddugoda, M., Huschtscha, L., Reddel, R., Chenevix-Trench, G., Toso, M., Kench, J.G., Horvath, L.G., et al. (2018). Guidelines for whole genome bisulphite sequencing of intact and FFPET DNA on the Illumina HiSeq X Ten. Epigenetics Chromatin 11, 24. 10.1186/s13072-018-0194-0.

42. Ming, X., Zhang, Z., Zou, Z., Lv, C., Dong, Q., He, Q., Yi, Y., Li, Y., Wang, H., and Zhu, B. (2020). Kinetics and mechanisms of mitotic inheritance of DNA methylation and their roles in aging-associated methylome deterioration. Cell Res. 30, 980–996. 10.1038/s41422-020-0359-9.

43. Iorio, F., Knijnenburg, T.A., Vis, D.J., Bignell, G.R., Menden, M.P., Schubert, M., Aben, N., Gonçalves, E., Barthorpe, S., Lightfoot, H., et al. (2016). A Landscape of Pharmacogenomic Interactions in Cancer. Cell 166, 740–754. 10.1016/j.cell.2016.06.017.

44. Charlton, J., Downing, T.L., Smith, Z.D., Gu, H., Clement, K., Pop, R., Akopian, V., Klages, S., Santos, D.P., Tsankov, A.M., et al. (2018). Global delay in nascent strand DNA methylation. Nat. Struct. Mol. Biol. 25, 327–332. 10.1038/s41594-018-0046-4.

45. Ren, H., Taylor, R.B., Downing, T.L., and Read, E.L. (2022). Locally correlated kinetics of post-replication DNA methylation reveals processivity and region specificity in DNA methylation maintenance. J. R. Soc. Interface 19, 20220415. 10.1098/rsif.2022.0415.

46. Vaughan, R.M., Dickson, B.M., Whelihan, M.F., Johnstone, A.L., Cornett, E.M., Cheek, M.A., Ausherman, C.A., Cowles, M.W., Sun, Z.-W., and Rothbart, S.B. (2018). Chromatin structure and its chemical modifications regulate the ubiquitin ligase substrate selectivity of UHRF1. Proc. Natl. Acad. Sci. 115, 8775–8780. 10.1073/pnas.1806373115.

47. Karg, E., Smets, M., Ryan, J., Forné, I., Qin, W., Mulholland, C.B., Kalideris, G., Imhof, A., Bultmann, S., and Leonhardt, H. (2017). Ubiquitome Analysis Reveals PCNA-Associated Factor 15 (PAF15) as a Specific Ubiquitination Target of UHRF1 in Embryonic Stem Cells. J. Mol. Biol. 429, 3814–3824. 10.1016/j.jmb.2017.10.014.

48. Nishiyama, A., Mulholland, C.B., Bultmann, S., Kori, S., Endo, A., Saeki, Y., Qin, W., Trummer, C., Chiba, Y., Yokoyama, H., et al. (2020). Two distinct modes of DNMT1 recruitment ensure stable maintenance DNA methylation. Nat. Commun. 11, 1222. 10.1038/s41467-020-15006-4.

49. Kong, X., Chen, J., Xie, W., Brown, S.M., Cai, Y., Wu, K., Fan, D., Nie, Y., Yegnasubramanian, S., Tiedemann, R.L., et al. (2019). Defining UHRF1 Domains that Support Maintenance of Human Colon Cancer DNA Methylation and Oncogenic Properties. Cancer Cell 35, 633–648.e7. 10.1016/j.ccell.2019.03.003.

50. Vaughan, R.M., Kupai, A., Foley, C.A., Sagum, C.A., Tibben, B.M., Eden, H.E., Tiedemann, R.L., Berryhill, C.A., Patel, V., Shaw, K.M., et al. (2020). The histone and non-histone methyllysine reader activities of the UHRF1 tandem Tudor domain are dispensable for the propagation of aberrant DNA methylation patterning in cancer cells. Epigenetics Chromatin 13, 44. 10.1186/s13072-020-00366-4.

51. Zhao, Q., Zhang, J., Chen, R., Wang, L., Li, B., Cheng, H., Duan, X., Zhu, H., Wei, W., Li, J., et al. (2016). Dissecting the precise role of H3K9 methylation in crosstalk with DNA maintenance methylation in mammals. Nat. Commun. 7, 12464. 10.1038/ncomms12464.

52. DaRosa, P.A., Harrison, J.S., Zelter, A., Davis, T.N., Brzovic, P., Kuhlman, B., and Klevit, R.E. (2018). A Bifunctional Role for the UHRF1 UBL Domain in the Control of Hemi-methylated DNA-Dependent Histone Ubiquitylation. Mol. Cell 72, 753–765.e6. 10.1016/j.molcel.2018.09.029.

53. Foster, B.M., Stolz, P., Mulholland, C.B., Montoya, A., Kramer, H., Bultmann, S., and Bartke, T. (2018). Critical Role of the UBL Domain in Stimulating the E3 Ubiquitin Ligase Activity of UHRF1 toward Chromatin. Mol. Cell 72, 739–752.e9. 10.1016/j.molcel.2018.09.028.

54. Jenkins, Y., Markovtsov, V., Lang, W., Sharma, P., Pearsall, D., Warner, J., Franci, C., Huang, B., Huang, J., Yam, G.C., et al. (2005). Critical Role of the Ubiquitin Ligase Activity of UHRF1, a Nuclear RING Finger Protein, in Tumor Cell Growth. Mol. Biol. Cell 16, 5621–5629. 10.1091/mbc.e05-03-0194.

55. Xu, C., and Corces, V.G. (2018). Nascent DNA methylome mapping reveals inheritance of hemimethylation at CTCF/cohesin sites. Science 359, 1166–1170. 10.1126/science.aan5480.

56. Stewart-Morgan, K.R., Requena, C.E., Flury, V., Du, Q., Heckhausen, Z., Hajkova, P., and Groth, A. (2023). Quantifying propagation of DNA methylation and hydroxymethylation with iDEMS. Nat. Cell Biol., 1–11. 10.1038/s41556-022-01048-x.

57. Busto-Moner, L., Morival, J., Ren, H., Fahim, A., Reitz, Z., Downing, T.L., and Read, E.L. (2020). Stochastic modeling reveals kinetic heterogeneity in post-replication DNA methylation. PLOS Comput. Biol. 16, e1007195. 10.1371/journal.pcbi.1007195.

58. Yuan, T., Jiao, Y., Jong, S. de, Ophoff, R.A., Beck, S., and Teschendorff, A.E. (2015). An Integrative Multi-scale Analysis of the Dynamic DNA Methylation Landscape in Aging. PLOS Genet. 11, e1004996. 10.1371/journal.pgen.1004996.

59. Heyn, H., Li, N., Ferreira, H.J., Moran, S., Pisano, D.G., Gomez, A., Diez, J., Sanchez-Mut, J.V., Setien, F., Carmona, F.J., et al. (2012). Distinct DNA methylomes of newborns and centenarians. Proc. Natl. Acad. Sci. 109, 10522–10527. 10.1073/pnas.1120658109.

60. Higham, J., Kerr, L., Zhang, Q., Walker, R.M., Harris, S.E., Howard, D.M., Hawkins, E.L., Sandu, A.-L., Steele, J.D., Waiter, G.D., et al. (2022). Local CpG density affects the trajectory and variance of age-associated DNA methylation changes. Genome Biol. 23, 216. 10.1186/s13059-022-02787-8.

61. Song, J., Rechkoblit, O., Bestor, T.H., and Patel, D.J. (2011). Structure of DNMT1-DNA Complex Reveals a Role for Autoinhibition in Maintenance DNA Methylation. Science 331, 1036–1040. 10.1126/science.1195380.

62. Song, J., Teplova, M., Ishibe-Murakami, S., and Patel, D.J. (2012). Structure-Based Mechanistic Insights into DNMT1-Mediated Maintenance DNA Methylation. Science 335, 709–712. 10.1126/science.1214453.

63. Bashtrykov, P., Rajavelu, A., Hackner, B., Ragozin, S., Carell, T., and Jeltsch, A. (2014). Targeted Mutagenesis Results in an Activation of DNA Methyltransferase 1 and Confirms an Autoinhibitory Role of its RFTS Domain. ChemBioChem 15, 743–748. 10.1002/cbic.201300740.

64. Kikuchi, A., Onoda, H., Yamaguchi, K., Kori, S., Matsuzawa, S., Chiba, Y., Tanimoto, S., Yoshimi, S., Sato, H., Yamagata, A., et al. (2022). Structural basis for activation of DNMT1. Nat. Commun. 13, 7130. 10.1038/s41467-022-34779-4.

65. Zhang, X., Li, B., Rezaeian, A.H., Xu, X., Chou, P.-C., Jin, G., Han, F., Pan, B.-S., Wang, C.-Y., Long, J., et al. (2017). H3 ubiquitination by NEDD4 regulates H3 acetylation and tumorigenesis. Nat. Commun. 8, 14799. 10.1038/ncomms14799.

66. Kim, C.R., Noda, T., Kim, H., Kim, G., Park, S., Na, Y., Oura, S., Shimada, K., Bang, I., Ahn, J.-Y., et al. (2020). PHF7 Modulates BRDT Stability and Histone-to-Protamine Exchange during Spermiogenesis. Cell Rep. 32, 107950. 10.1016/j.celrep.2020.107950.

67. Lee, H.S., Bang, I., You, J., Jeong, T.-K., Kim, C.R., Hwang, M., Kim, J.-S., Baek, S.H., Song, J.-J., and Choi, H.-J. (2023). Molecular basis for PHF7-mediated ubiquitination of histone H3. Genes Dev. 37, 984–997. 10.1101/gad.350989.123.

68. Wang, H., Zhai, L., Xu, J., Joo, H.-Y., Jackson, S., Erdjument-Bromage, H., Tempst, P., Xiong, Y., and Zhang, Y. (2006). Histone H3 and H4 Ubiquitylation by the CUL4-DDB-ROC1 Ubiquitin Ligase Facilitates Cellular Response to DNA Damage. Mol. Cell 22, 383–394. 10.1016/j.molcel.2006.03.035.

69. Vaughan, R.M., Kupai, A., and Rothbart, S.B. (2021). Chromatin Regulation through Ubiquitin and Ubiquitin-like Histone Modifications. Trends Biochem. Sci. 46, 258–269. 10.1016/j.tibs.2020.11.005.

70. Wiederschain, D., Susan, W., Chen, L., Loo, A., Yang, G., Huang, A., Chen, Y., Caponigro, G., Yao, Y., Lengauer, C., et al. (2009). Single-vector inducible lentiviral RNAi system for oncology target validation. Cell Cycle 8, 498–504. 10.4161/cc.8.3.7701.

71. Mungamuri, S.K., Qiao, R.F., Yao, S., Manfredi, J.J., Gu, W., and Aaronson, S.A. (2016). USP7 Enforces Heterochromatinization of p53 Target Promoters by Protecting SUV39H1 from MDM2-Mediated Degradation. Cell Rep. 14, 2528–2537. 10.1016/j.celrep.2016.02.049.

72. Chomiak, A.A., Tiedemann, R.L., Liu, Y., Kong, X., Cui, Y., Thurlow, K., Cornett, E.M., Topper, M.J., Baylin, S.B., and Rothbart, S.B. (2023). Select EZH2 inhibitors enhance the viral mimicry effects of DNMT inhibition through a mechanism involving calcium-calcineurin-NFAT signaling. Preprint at bioRxiv, 10.1101/2023.06.09.544393 10.1101/2023.06.09.544393.

73. Zhou, W., Triche, T.J., Jr, Laird, P.W., and Shen, H. (2018). SeSAMe: reducing artifactual detection of DNA methylation by Infinium BeadChips in genomic deletions. Nucleic Acids Res. 46, e123. 10.1093/nar/gky691.

74. Gentleman, R.C., Carey, V.J., Bates, D.M., Bolstad, B., Dettling, M., Dudoit, S., Ellis, B., Gautier, L., Ge, Y., Gentry, J., et al. (2004). Bioconductor: open software development for computational biology and bioinformatics. Genome Biol. 5, R80. 10.1186/gb-2004-5-10-r80.

75. Huber, W., Carey, V.J., Gentleman, R., Anders, S., Carlson, M., Carvalho, B.S., Bravo, H.C., Davis, S., Gatto, L., Girke, T., et al. (2015). Orchestrating high-throughput genomic analysis with Bioconductor. Nat. Methods 12, 115–121. 10.1038/nmeth.3252.

76. Burger, L., Gaidatzis, D., Schübeler, D., and Stadler, M.B. (2013). Identification of active regulatory regions from DNA methylation data. Nucleic Acids Res. 41, e155–e155. 10.1093/nar/gkt599.

77. Heinz, S., Benner, C., Spann, N., Bertolino, E., Lin, Y.C., Laslo, P., Cheng, J.X., Murre, C., Singh, H., and Glass, C.K. (2010). Simple Combinations of Lineage-Determining Transcription Factors Prime *cis*-Regulatory Elements Required for Macrophage and B Cell Identities. Mol. Cell 38, 576–589. 10.1016/j.molcel.2010.05.004.

78. Zhao, P.A., Sasaki, T., and Gilbert, D.M. (2020). High-resolution Repli-Seq defines the temporal choreography of initiation, elongation and termination of replication in mammalian cells. Genome Biol. 21, 76. 10.1186/s13059-020-01983-8.

79. Faust, G.G., and Hall, I.M. (2014). SAMBLASTER: fast duplicate marking and structural variant read extraction. Bioinformatics 30, 2503–2505. 10.1093/bioinformatics/btu314.

80. Ramírez, F., Dündar, F., Diehl, S., Grüning, B.A., and Manke, T. (2014). deepTools: a flexible platform for exploring deep-sequencing data. Nucleic Acids Res. 42, W187–W191. 10.1093/nar/gku365.

81. Lund, E., Oldenburg, A.R., and Collas, P. (2014). Enriched domain detector: a program for detection of wide genomic enrichment domains robust against local variations. Nucleic Acids Res. 42, e92. 10.1093/nar/gku324.

82. Quinlan, A.R., and Hall, I.M. (2010). BEDTools: a flexible suite of utilities for comparing genomic features. Bioinformatics 26, 841–842. 10.1093/bioinformatics/btq033.

83. Vaughan, R.M., Dickson, B.M., Cornett, E.M., Harrison, J.S., Kuhlman, B., and Rothbart, S.B. (2018). Comparative biochemical analysis of UHRF proteins reveals molecular mechanisms that uncouple UHRF2 from DNA methylation maintenance. Nucleic Acids Res. 46, 4405–4416. 10.1093/nar/gky151.

84. Hann, Z.S., Ji, C., Olsen, S.K., Lu, X., Lux, M.C., Tan, D.S., and Lima, C.D. (2019). Structural basis for adenylation and thioester bond formation in the ubiquitin E1. Proc. Natl. Acad. Sci. 116, 15475–15484. 10.1073/pnas.1905488116.

