## Supplemental Figures for "UHRF1 ubiquitin ligase activity supports the maintenance of low-density CpG methylation"

SUPPLEMENTARY INFORMATION

### Figure S1. Tiedemann et al.

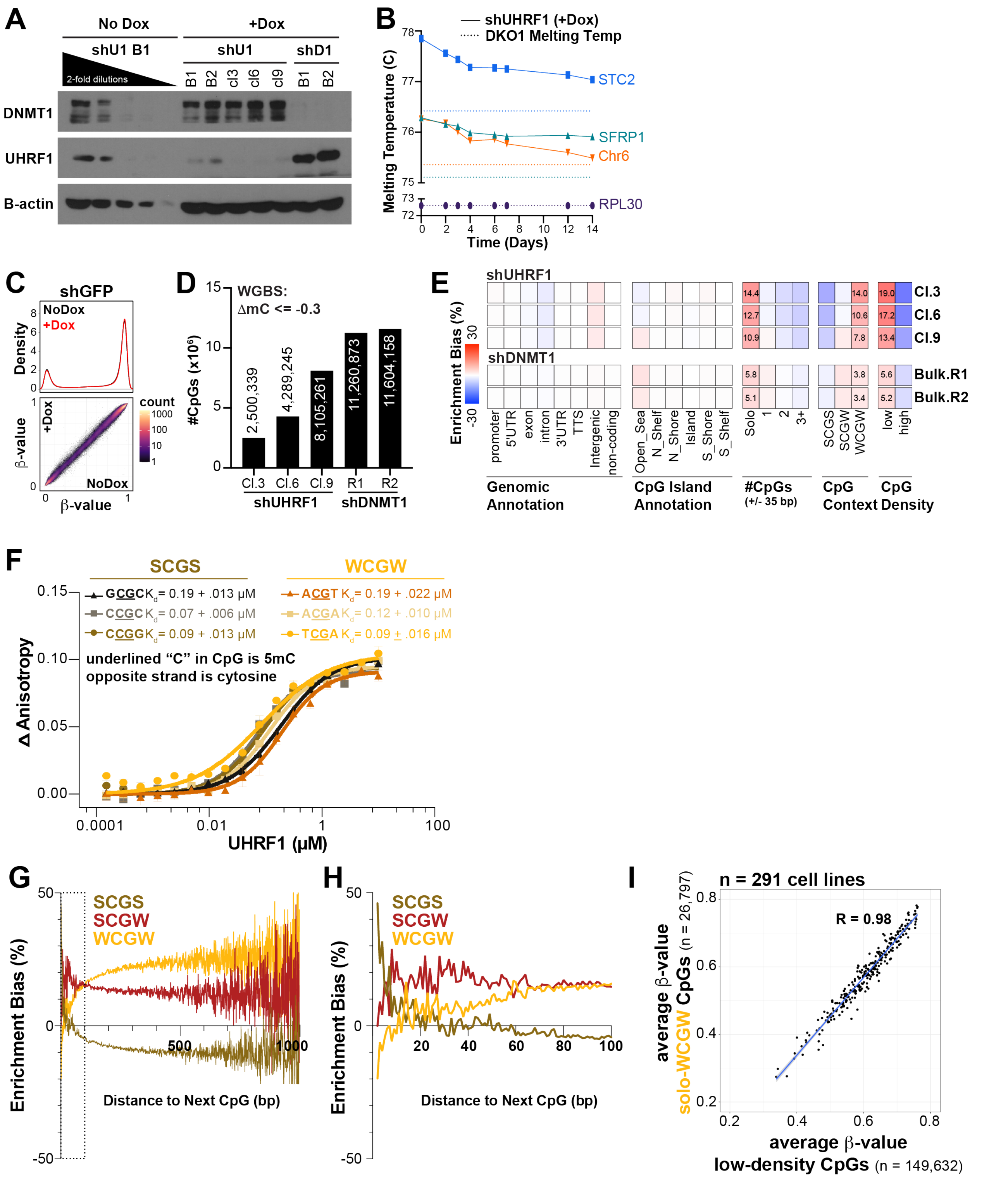

**Figure S1. Validation of model and additional analysis of Whole Genome Bisulfite Sequencing (WGBS) (A)** Western blot analysis of UHRF1/DNMT1 shRNA knockdown in bulk (B) and clonal (cl) populations of engineered stable doxycycline-inducible shRNA HCT116 cells. For comparison of knockdown efficiency among samples, the lysate from the NoDox UHRF1 bulk sample is titrated in 2-fold dilutions starting from 10 ug of protein. **(B)** High Resolution Melt (HRM) analysis of amplicons from different genomic regions across collected time-points in the shUHRF1 cl.6 HCT116 cells. DNA methylation levels are inferred from the melting temperature of the amplicon where higher melting temperatures indicate higher levels of DNA methylation and lower melting temperatures indicate lower levels of DNA methylation. DNA from HCT116 DKO1 cells is used as a positive control for loss of DNA methylation. STC2 = gene body region, SFRP1 = promoter region, Chr6 = pericentric heterochromatin region on Chr6, RPL30 = promoter, unmethylated control. **(C)** DNA methylation distributions of EPIC array probes from doxycycline (dox)-inducible shGFP HCT116 cell lines without (black) and with (red) dox treatment (10 ng/mL) for 14 days. **Top panel:** Density plots for CpG probe distribution across DNA methylation levels ($\beta$-value: 0 (unmethylated) to 1 (methylated)). **Bottom panel:** Density scatterplots demonstrating density of probes and DNA methylation level in Baseline (x-axis) and knockdown (y-axis) methylomes. **(D)** Bar graph for number of differentially methylated CpGs (WGBS) for each knockdown experiment ($mC_{Baseline} \geq 0.85$; $\Delta mC_{Knockdown} \leq -0.3$). **(E)** Hypergeometric analysis of significantly hypomethylated CpGs for each sample from **S1E.** Red indicates significant overrepresentation for hypomethylation of the feature and blue indicates significant underrepresentation. Enrichment bias values are provided for the most significant positive enrichments. #CpGs indicates the number of CpGs $\pm$ 35 bp upstream and downstream of the hypomethylated CpG. CpG Context represents the -1/+1 position nucleotide (S = C or G; W = A or T) flanking the hypomethylated CpG. CpG density is determined by the #bps to the next CpG (either upstream or downstream). Low density $\geq$ 20 bp, High Density < 20 bp. **(F)** Fluorescence polarization of full-length UHRF1 with a fluorescent 12 bp DNA probe differing at nucleotides (-/+ 1 bp) flanking the centrally-located CpG. **(G)** Hypergeometric analysis of all CpGs in the genome by flanking nucleotide context and CpG binning by distance to the next CpG (bp). Positive enrichment bias indicates overrepresentation, negative enrichment bias indicates underrepresentation. Dotted box indicates zoomed in enrichment bias from 0 to 100 bp distance presented in **S1H**. **(H)** Enrichment bias results from dotted box in **S1G. (I)** Scatterplot of calculated average $\beta$-value for low-density CpGs (Distance to Next CpG $\geq$ 20 bp, n = 149,632) versus average $\beta$-value for solo-WCGW (n = 26,797) from 450K array analysis of 291 cell lines (GSE68379). **See also Figure 1.**

#### Figure S2. Tiedemann et al.

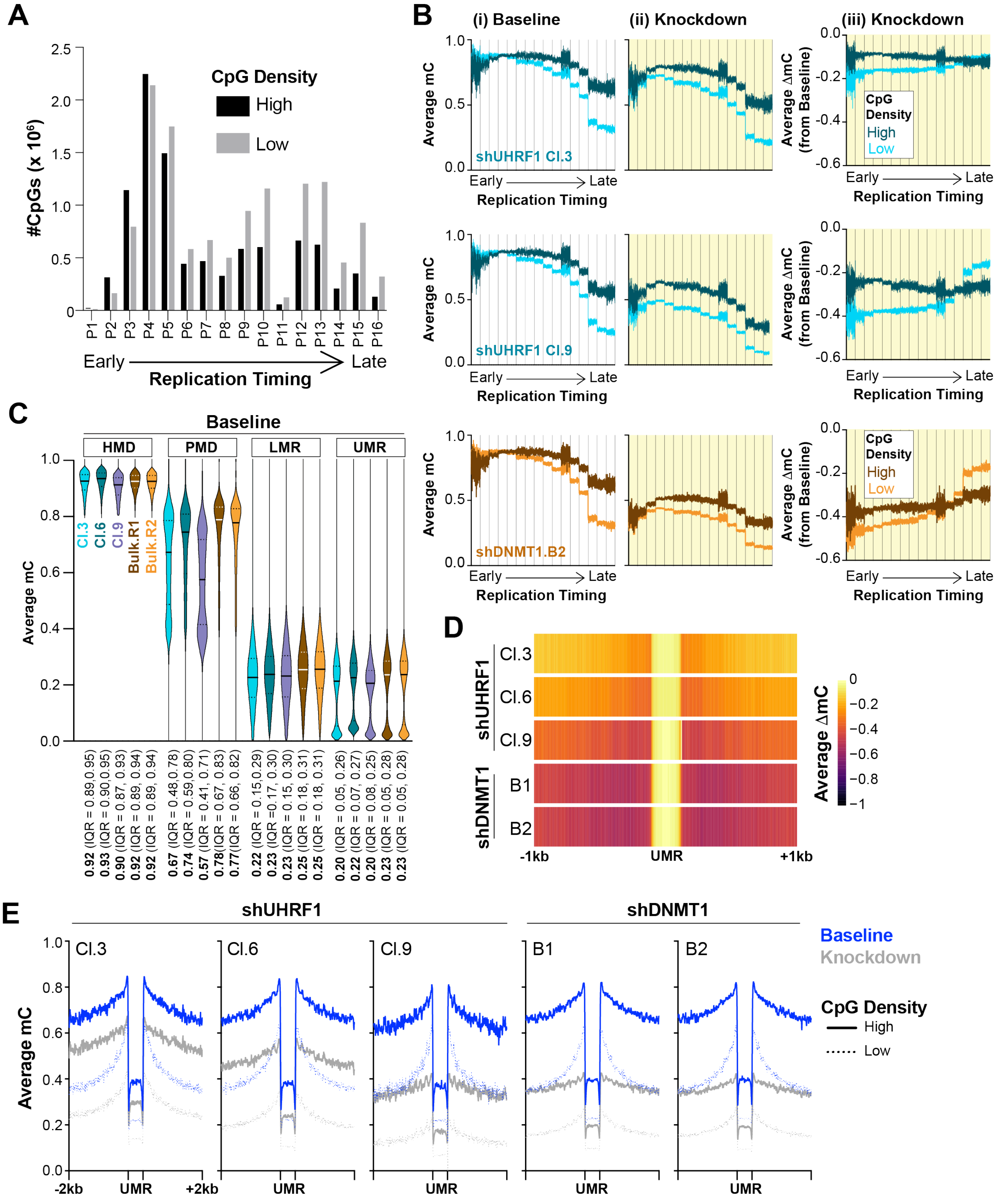

**Figure S2. Replication timing integration of additional samples and analysis of expanding UMRs** **(A)** Distribution of high- and low-density CpGs across replication timing phases. Note low-density CpGs are enriched in late-replicating regions. **(B)** Average DNA methylation (mC) of high (dark color) and low (light color) density CpGs in (i) Baseline and (ii) Knockdown phases and the (iii) average change in methylation ($\Delta mC_{Knockdown}$ across each replication timing phase (16-phase) in HCT116 cells. Replication timing phases are assigned in 50kb windows genome-wide, and the average DNA methylation (from WGBS) is calculated in 100 bp bins from the start of the 50kb window to the end for each timing phase. **(C)** Average DNA methylation (mC) distributions among called Methylation Domains in Baseline Methylomes. Values presented on x-axis indicate median(Interquartile Range (IQR) for 25-75% of the data). **(D)** Heatmaps for loss of DNA methylation adjacent to UMRs within PMDs. **(E)** Average DNA methylation profiles (mC) of Baseline (blue) and Knockdown (gray) methylomes at UMRs present in Baseline PMDs. High-density CpGs are denoted by solid lines and low-density CpGs with dotted lines across shUHRF1 and shDNMT1 samples (WGBS). **See also Figure 2.**

### Figure S3. Tiedemann et al.

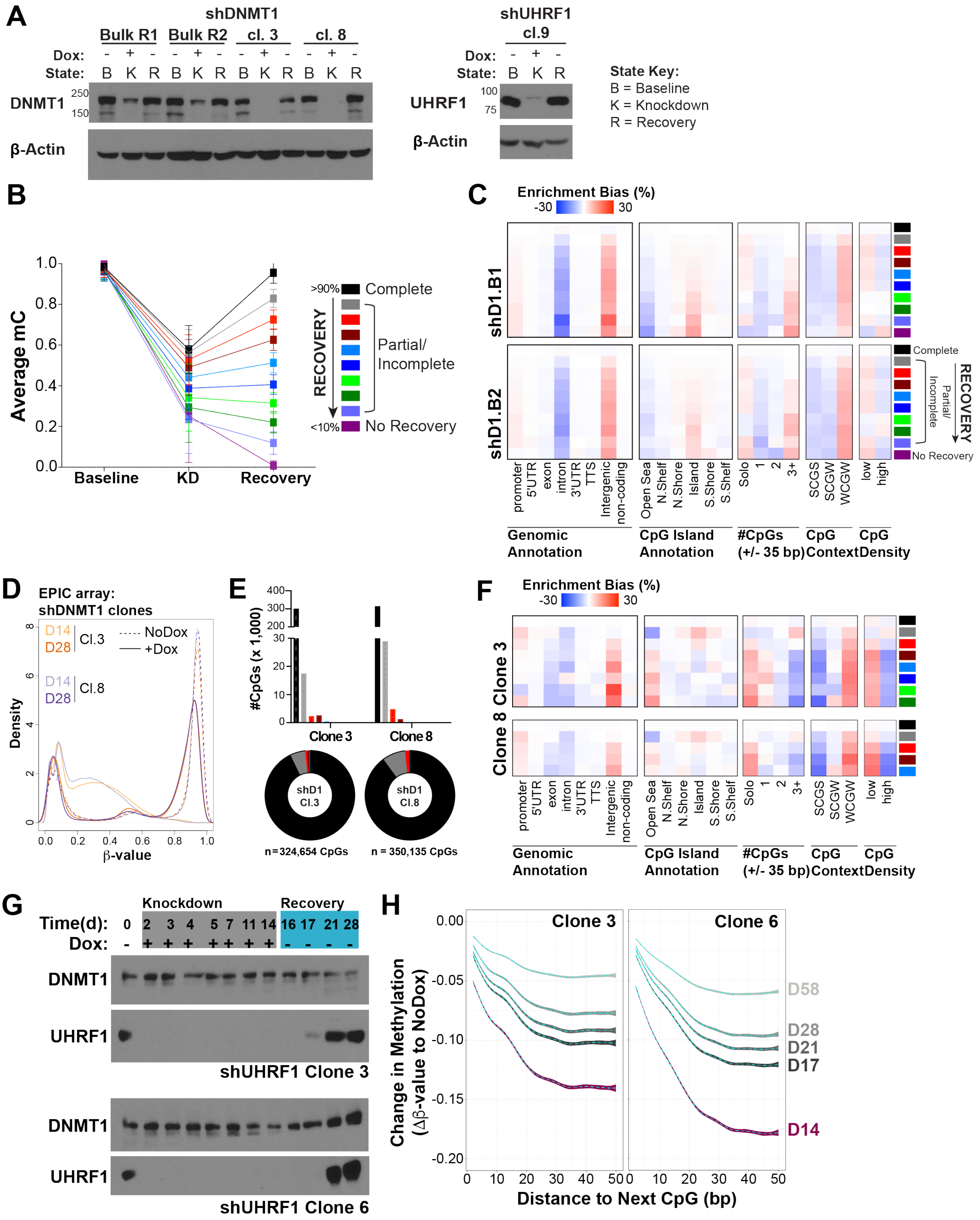

**Figure S3. Recovery validation and analysis (A)** Western blot analysis of DNMT1 (Bulk and clonal populations) and UHRF1 (Cl.9, note: Cl.3,6 presented in **Figure S1G**) recovery protein expression following two weeks of dox removal. **(B)** Recovery analysis for significantly hypomethylated CpGs (from knockdown) in shUHRF1 cl.6 (WGBS). DNA methylation recovery is determined by calculating the $\Delta$mC of Day 28 samples (recovery) to the respective Baseline (NoDox) samples. DNA methylation recovery measurements were then binned into three groups, where CpGs with recovery values ($|\Delta mC_{Recovery}|$*100) < 10% are considered not recovered, > 90% are considered complete recovery, and between 10 and 90% are considered incomplete or partial recovery. **(C)** Hypergeometric analysis of DNA methylation recovery for each shDNMT1 sample (WGBS) from **Figure 3A.** Red indicates significant overrepresentation for recovery of the feature (and blue indicates significant underrepresentation. **(D)** DNA methylation distributions of EPIC array probes from dox-inducible shDNMT1 clonal HCT116 cell lines for Baseline, Knockdown, and Recovery phases. **(E)** Distribution of recovery categories among shDNMT1 clonal samples (EPIC). Scale from **S3B** applies. **(F)** Enrichment bias analysis of different recovery phases for clonal shDNMT1 samples. (NOTE: If a recovery phase had less than 30 CpGs, the phase was omitted due to low sampling number for hypergeometric testing). **(G)** Western blot analysis of UHRF1 recovery protein expression following two weeks of dox removal in Cl.3 and Cl.6. **(H)** Average loss of DNA methylation for all highly methylated CpGs ($\beta$-value$_{Baseline}$ $\geq$ 0.85) across 'Distance to the Next CpG' binning across recovery time points. Dotted line indicates the average Db-value as a function of distance, colored boundaries indicate 95% confidence intervals. **See also Figure 3.**

Fig. S4 Tiedemann et al.

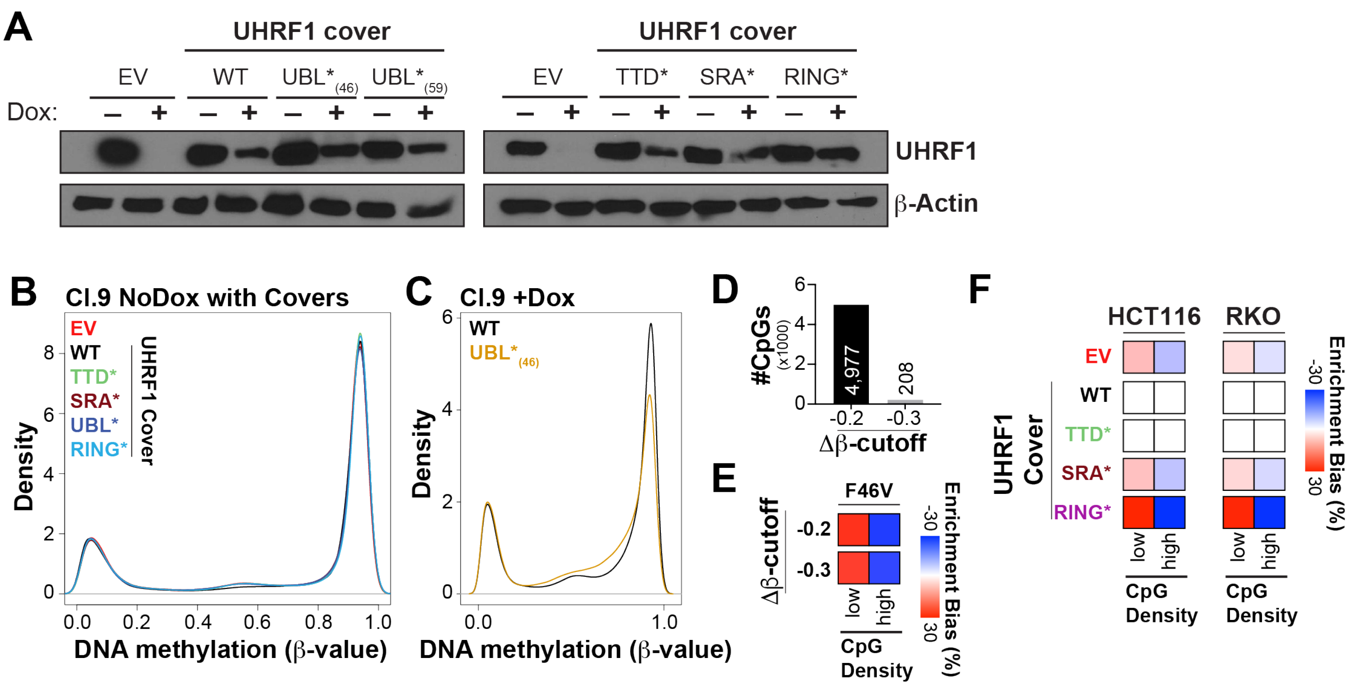


**Figure S4. UHRF1 transgene cover additional analysis (A)** Western blot analysis of UHRF1 transgene cover expression over the shUHRF1 Cl.9 HCT116 background. Cells were treated with doxycycline (10 ng/ul) for 14 days to induce knockdown of endogenous UHRF1 and determine methylation maintenance in the presence of the UHRF1 transgene cover. **(B)** DNA methylation distributions of the NoDox samples with UHRF1 transgene covers (prior to any doxycycline exposure) to demonstrate no changes in the Baseline methylome. **(C)** DNA methylation distribution of the WT UHRF1 cover (+Dox) and UBL*$_{(F46V)}$ UHRF1 cover (+Dox). **(D)** Bar graph of number of hypomethylated CpGs (EPIC array) for UBL*$_{(F46V)}$ UHRF1 cover (β-value$_{WT_cover}$ ≥ 0.85; varying Db-value$_{UBL*(F46V)}$ cutoffs). **(E)** Hypergeometric analysis of hypomethylated CpGs from UBL*$_{(F46V)}$ UHRF1 cover experiment for CpG density. **(H)** Hypergeometic analysis of hypomethylated CpGs from Kong et. al. 2018 UHRF1 cover experimetns in HCT116 and RKO colorectal cancer cells. **See also Figure 4.**

**Fig. S5 Tiedemann et al.**

**A**

| DNMT1 RFTS | Nucleosome | | |
|---|---|---|---|
| | H3K14ub | H3K18ub | H3K23ub |
| WT | 10 nM +/- 0.3 | 10 nM +/- 0.2 | 44 nM +/- 2.1 |
| UIM1* | 35 nM +/- 0.5 | 66 nM +/- 0.9 | 140 nM +/- 7.6 |
| UIM2* | n.m. | 891 nM +/- 25 | n.m. |
| dblUIM* | n.m. | n.m. | n.m. |

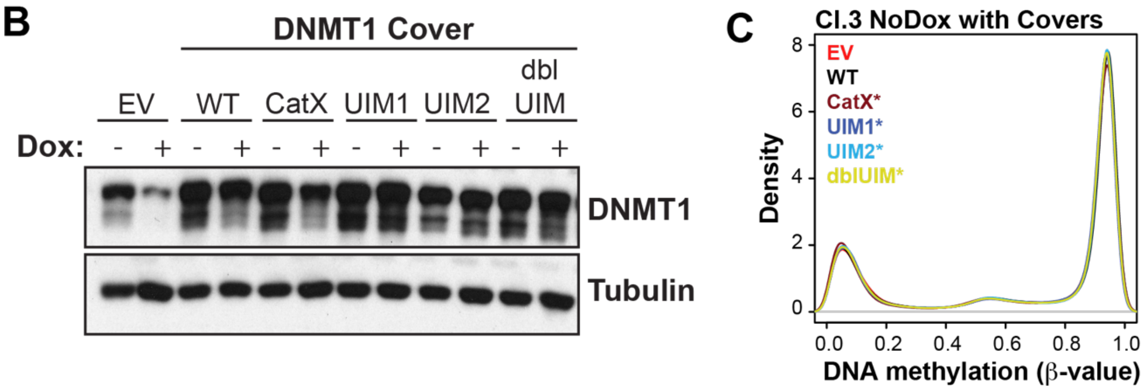


**Figure S5. DNMT1 transgene cover additional analysis (A)** Kd values (in nM) measured by Alpha screen for interactions between recombinant DNMT1 RFTS domains (wild type and mutants) and H3 ubiquitinated nucleosomes (K14ub, K18ub, K23ub). **(B)** Western blot analysis of DNMT1 transgene cover expression over the shDNMT1 Cl.3 HCT116 background. Cells were treated with doxycycline (10 ng/ul) for 14 days to induce knockdown of endogenous DNMT1 and determine methylation maintenance in the presence of the DNMT1 transgene covers. **(C)** DNA methylation distributions of the NoDox samples with DNMT1 transgene covers (prior to any doxycycline exposure) to demonstrate no changes in the Baseline methylome. **See also Figure 5.**
